# Pharmacological tuning of microRNAs in FAP-derived Extracellular Vesicles by HDAC inhibitors promotes regeneration and reduces fibrosis in dystrophic muscles

**DOI:** 10.1101/2020.02.15.947325

**Authors:** M. Sandonà, S. Consalvi, L. Tucciarone, M. De Bardi, M. Scimeca, D. Angelini, V. Buffa, A. D’Amico, E. Bertini, S. Cazzaniga, P. Bettica, M. Bouché, A. Bongiovanni, P.L. Puri, V. Saccone

## Abstract

Functional interactions between cellular components of the muscle stem cell (MuSC) niche regulate the regenerative ability of skeletal muscles in physiological and pathological conditions; however, the identity of the mediators of these interactions remains largely unknown. We show here that fibro-adipogenic progenitor (FAP)-derived Extracellular Vesicles (EVs) mediate microRNA transfer to MuSCs, and that exposure of dystrophic FAPs to HDAC inhibitors (HDACi) increases the intra-EV levels of a subset of microRNAs (miRs), which cooperatively target biological processes of therapeutic interest, including regeneration, fibrosis and inflammation. In particular, we found that increased levels of miR206 in EVs released from FAPs of muscles from Duchenne dystrophic patients or mice (mdx) exposed to HDACi were associated with enhanced regeneration and inhibition of fibrosis of dystrophic muscles. Consistently, EVs from HDACi-treated dystrophic FAPs could stimulated MuSC activation and expansion *ex vivo,* and promoted regeneration, while inhibiting fibrosis and inflammation of dystrophic muscles, upon intramuscular transplantation, *in vivo.* These data reveal a potential for pharmacological modulation of FAP-derived EV’s content as novel strategy for focal therapeutic interventions in Duchenne Muscular Dystrophy (DMD) and possibly other muscular diseases.

**Brief Summary:** Extracellular Vesicles from HDACi-treated dystrophic FAPs promote regeneration, while inhibiting fibrosis and inflammation of dystrophic muscles

## Introduction

Emerging evidence indicates that reciprocal interactions between distinct cellular components of the regeneration machinery generate either productive or hostile environment for regeneration of dystrophic muscles (*1–3*). While muscle stem (satellite) cells(*4*) (therein indicated as MuSCs) are the direct effectors of muscle repair, a variety of other cell types contribute to muscle regeneration by spatially and temporally coordinating the activity of MuSCs(*5, 6*). These cells comprise components of the inflammatory infiltrate, including macrophages and eosinophils(*7, 8*), and a heterogeneous population of muscle-derived interstitial cells referred to as fibro-adipogenic progenitors (FAPs)(*9–11*). Disruption of this network compromises the integrity of MuSC niche and has been associated with the progression of many chronic muscular disorders (i.e. muscular dystrophies) and age-related decline in muscle mass and repair(*12, 13*).

Upon myofiber damage, FAP accumulation is preceded by the appearance of the inflammatory infiltrate and is followed by MuSC activation(*14, 15*). This temporal pattern suggests a key role for FAPs in converting inflammatory cues into signals that regulate MuSC activity, and implies reciprocal communications between these cell types, through the exchange of soluble mediators, which are largely unknown. Deciphering the molecular and functional identity of these mediators might reveal selective targets for interventions aimed at restoring functional interactions between cellular components of the skeletal muscle environment during maladaptive repair in chronic muscular disorders.

Previous studies identified FAPs as key cellular targets of histone deacetylase inhibitors (HDACi) – a pharmacological intervention that counters DMD progression by promoting compensatory regeneration, while inhibiting fibro-adipogenic degeneration both in pre-clinical studies(*16–22*) and clinical trials(*23*). While these studies indicate the pharmacological potential of HDACi to restore the regenerative environment in dystrophic muscles, by recovering physiological functional interactions between FAPs and other cellular components(*24*), the signals that mediate HDACi ability to restore physiological interactions between FAPs and other cell types in DMD muscles remain largely unknown. Here, we report the identification and functional characterization of HDACi-induced extracellular vesicles (EVs) that mediate FAP’s ability to promote MuSCs activation and differentiation, as well as to inhibit muscle fibrosis and inflammation, in DMD muscles.

## Results

### EVs mediate FAP ability to promote MuSCs activation and differentiation upon exposure to HDACi

The HDACi Givinostat (Giv) is the first epigenetic drug that has been used in a clinical trial with DMD boys(*23*). Histological analysis of muscles from patients treated with Giv showed increased regeneration, with a concomitant decrease of fibrosis and fat infiltration, similar to those observed in pre-clinical studies with mdx mice (the mouse model of DMD) treated with pan HDACi, such as Giv and Trichostatin A (TSA), which exhibit equivalent activities(*19*). In addition to the histological analysis, we also obtained access to representative biopsies from 2 DMD patients, before and after Giv treatment, from which we isolated by FACS two populations of our main interest – human MuSCs (CD11b neg, CD31 neg, CD45 neg, CD56/NCAM pos) and a population enriched in presumptive human FAPs (CD11b neg, CD31 neg, CD45neg, CD56/NCAMneg). Fig. S1A shows that patient-derived human MuSCs (hMuSCs) could form myotubes when cultured *in vitro*; likewise, the cell population enriched in human FAPs (ehFAPs) could differentiate into adipocytes *in vitro* when cultured in adipogenic medium. The availability of these patient-derived cells provides an unprecedented opportunity to evaluate the effect of HDACi on MuSC-FAPs interactions *ex vivo* and *in vivo*. *Ex vivo* experiments were performed by transwell co-culture of hMuSCs and ehFAPs from patients before the beginning of the trial. In both patients, co-culture with ehFAPs slightly decreased hMuSC ability to differentiate into multinucleated myotubes (Fig. 1A-B). This is in contrast with the reported ability of FAPs isolated from 2 month old mdx mice to promote differentiation of MuSCs from the same animal(*18*). The discrepancy between these results is likely accounted by differences of DMD progression between human patients and mouse models, with 8 to 11 years old patients enrolled in the clinical trial being at more advanced stages of disease progression, as also shown by the more pronounced fibrosis observed in their muscles, as compared to 2 months old mdx mice. Exposure of ehFAPs to Giv (ehFAPs Giv Vitro) prior to co-culture with hMuSCs could increase the formation of hMuSC-derived myotubes to an extent comparable to that observed in hMuSCs isolated from patients after one year of Giv treatment *in vivo* (hMuSCs Giv) (Fig. 1A,B). These data indicate that FAP’s ability to support MuSC differentiation is compromised in muscles of DMD patients, but can be recovered upon treatment with HDACi.

**Fig. 1:**
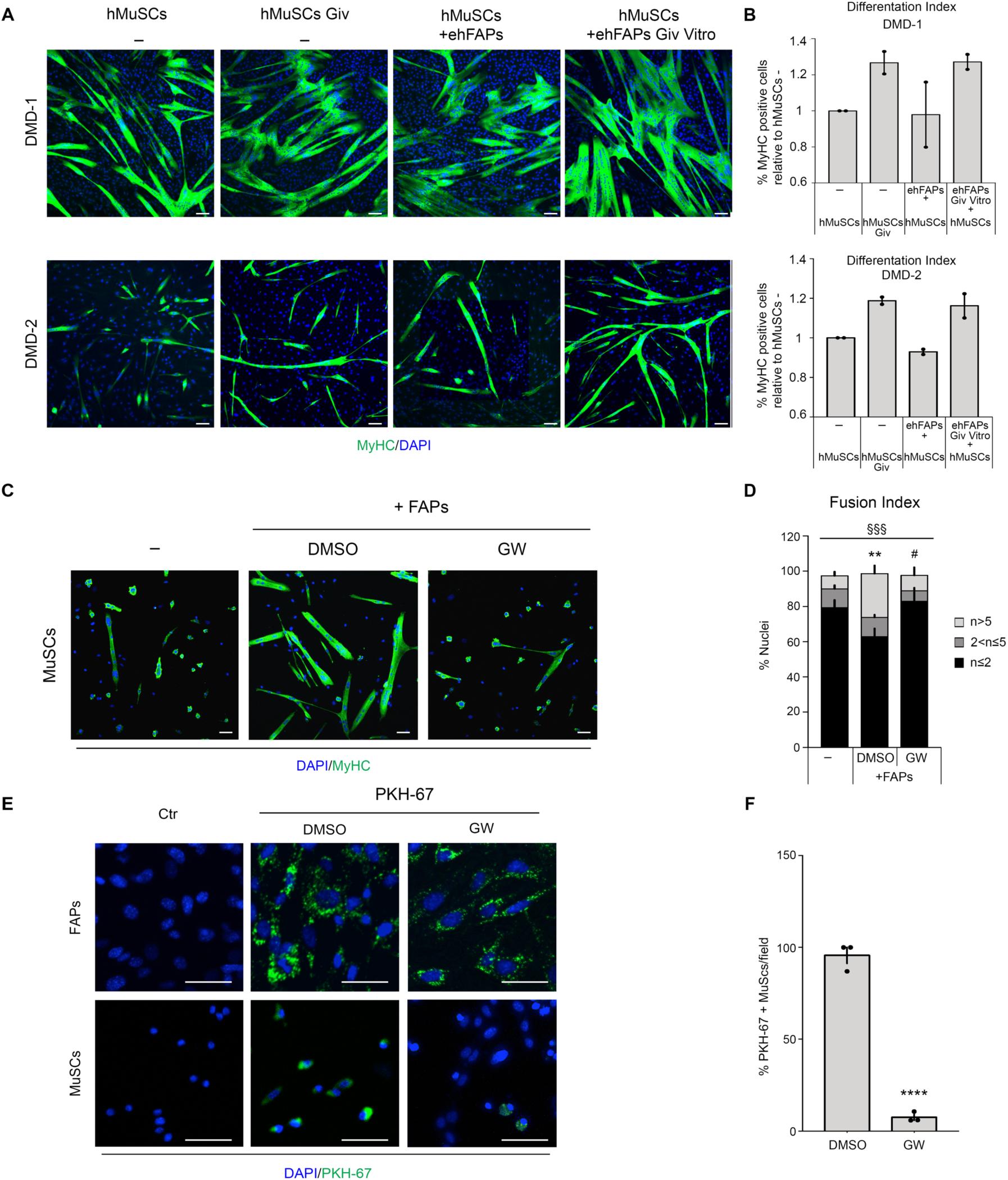
HDACi restore DMD FAP ability to support to MuSC differentiation into multinucleated myotubes. **(A)** Representative images showing the myogenic differentiation, as assessed by immunostaining for MyHC (green), of human MuSCs (hMuSCs) isolated from muscle biopsies of DMD patients, either before the beginning of the clinical trial (hMuSCs (-)) or after 1 year of treatment with Givinostat (hMuSCs Giv (-)). Cell were cultured alone or (only for hMuSCs-) in transwell co-culture with FAPs isolated from the same patients and exposed in vitro to Giv (+ehFAPs Giv Vitro) or control vehicle (+ehFAPs).The upper panels represent myogenic differentiation of DMD-1 patient; bottom panels represent myogenic differentiation of DMD-2 patient. Dose and timing of Giv treatment as described in Bettica et al., 2016 (23). (**B)** Graph showing the differentiation index of hMuSCs described in A. All data correspond to the average ± SD. (**C)** Representative images showing the myogenic differentiation of MuSCs isolated from 1.5 month old mdx mice, as assessed by immunostaining for MyHC (green). MuSCs were cultured alone (-) or in transwell co-culture with mdx FAPs, which were pretreated with DMSO or GW4869 (GW 10μM added to FAPs 30 minutes before starting the co-culture with MuSCs). Scale bar = 50 μm**. (D)** Graph showing the fusion index of MuSCs in the conditions described in C, (n=5). Star (*) indicates statistical analysis by Tukey test relative to MuSCs cultured alone (-), ** p < 0.01; Hash (#) indicates statistical analysis by Tukey test relative to MuSCs in co-cultured with FAPs not treated (DMSO), # p<0.05; § represents statistical analysis by 2Way Anova test; §§§ p < 0,001. (**E)** Representative images of PKH-67 (green) and DAPI (blue) staining in a transwell co-culture between MuSCs and FAPs isolated from 1.5 month old mdx mice. FAPs exposed to PKH-67 prior to the co-culture with MuSCs. Scale bar = 50 μm. (**F)** Graph showing the percentage of PKH-67 positive MuSCs after the co-culture with FAPs, (n=3). Star (*) indicates statistical analysis by t-test;**** p < 0.0001. Nuclei were counterstained with DAPI (blue). All data correspond to the average ± SEM.

To determine the identity of the extracellular mediators of the functional interactions between FAPs and MuSCs in DMD muscles, we performed transwell co-cultures of these cells isolated from the mouse model of Duchenne Muscular Dystrophy (DMD) – the mdx mice. FAPs were isolated from 1.5 month old mdx mice, and were co-cultured with MuSCs isolated from age-matched mdx mice. In transwell cultures, cells are separated by a membrane of 1 µm pore size that prevents direct cell contact, yet allows reciprocal transfer of soluble mediators (e.g. growth factors and cytokines as well as extracellular vesicles (EVs), such as exosomes) exchanged between co-cultured cells. Fig. 1 shows that FAPs enhanced the ability of co-cultured MuSCs to differentiate into multinucleated myotubes, as compared to MuSCs cultured alone (Fig. 1C-D), as previously reported (*18*). To determine the relative contribution of FAP-derived EVs, FAPs were treated with the neutral sphingomyelinase inhibitor GW4869, which selectively blocks exosome biogenesis (*25*) FAPs exposure to GW4869 (30 minutes before starting the co-culture with MuSCs) abrogated FAP’s ability to enhance MuSCs differentiation into multinucleated myotubes (Fig.1C and D). This evidence indicates that EVs are essential contributors of FAP-mediated support to MuSCs activity and implies that FAP-delivered EV/exosomes are transferred to MuSCs, whereby they influence MuSC biological properties.

We next isolated EVs from FAP’s supernatant and analyzed their size distribution by Dynamic Light Scattering (DLS) experiments. To this purpose we have compared two purification procedures; i) a commercially available Total Exosome Isolation^TM^ precipitation reagent (TEIR) and ii) the current gold standard isolation method, which is based on differential ultracentrifugations (UC)(*26, 27*) DLS analyses showed for both FAPs-derived vesicles preparations an average hydrodynamic diameter of about 150 nm that is consistent with the standard size of exosomes (fig. S1B)(*28*). DLS analysis was also complemented with scanning electron microscopy (SEM) analyses of fixed TEIR-isolated EVs that showed round-shaped FAP-derived EVs ranging from 100-150 nm in size (fig. S1C). The slight difference in size between SEM and DLS analyses can be explained by the fact that DLS measures the hydrodynamic diameter of native particles in dispersion and is inherently biased towards larger particles(*29*). Moreover, a western blot analysis of the protein content showed that presumptive exosome markers were abundantly expressed in EVs TEIR-isolated from FAP’s supernatant, including Alix, Hsp70, and to a less extent Flotillin1 and CD63 (fig. S1D). EVs preparation was cleared from contaminating cell organelles, as indicated by the absence of Calnexin, an ubiquitously expressed ER protein that was exclusively found in FAP’s whole cell fractions (WCL) (fig. S1D). While these data demonstrate that functionally active FAPs-derived EVs display typical features of mammalian exosomes(*30*), in this manuscript we will continue to refer to these nanoparticles as EVs.

To analyse the capacity of MuSCs to uptake FAP-derived EVs, we performed a transwell co-culture in which the potential cellular “source” of EVs – the FAPs - was transfected with the exosomal marker CD63 fused to GFP prior to transwell co-culture with the “acceptor” cells – the MuSCs. In this system, detection of GFP into MuSCs reveals the transfer of CD63-labelled EVs from FAPs to MuSCs. Fig. S1E shows that GFP signal was detected in FAPs after transfection with GFP-CD63, as well as in MuSCs co-cultured with FAPs previously transfected with GFP-CD63. By contrast, no GFP signal was detected in MuSCs co-cultured with FAPs previously transfected with a control plasmid (mock) (fig. S1E). To further demonstrate the transfer of EVs from FAPs to MuSCs, we incubated FAPs with the lipidic dye (PKH-67) prior to transwell co-culture with MuSCs. PKH-67 is incorporated into newly generated EVs; hence, it can be used to trace EV passage from FAPs to acceptor cells. Indeed, PKH-67 staining was invariably detected in FAPs exposed to the dye as well as in recipient co-cultured MuSCs. On the other hand, prior exposure of FAPs to GW4869 almost completely prevented the detection of the signal in MuSCs, (Fig. 1E, F), demonstrating that FAP-derived EVs are uptaken by MuSCs.

To evaluate whether FAP-derived EVs could deliver regulatory signals to MuSCs, we incubated MuSCs with EVs isolated from the supernatant of FAPs from 1.5 month old mdx mice (CTR). FAP-derived EVs increased the ability of MuSCs to form multinucleated myotubes (Fig. 2A and B – compare CTR to untreated-MuSCs (-)). To determine whether EVs mediate the abiliy of HDACi to enhance FAP’s support to MuSC differentiation, we incubated MuSCs with EVs that were purified from FAPs of mdx mice exposed for 15 days to TSA. These EVs further increase the formation of multinucleated myotubes from MuSCs, as compared to EVs from FAPs of CTR-treated mdx (Fig. 2A and B – compare TSA to CTR). We next sought to evaluate whether HDACi could directly promote release of pro-myogenic EVs, by exposing cultures of FAPs isolated from vehicle-treated mdx mice to TSA (TSA vitro). Fig. 2A and B shows that exposure of FAPs to TSA *ex vivo* led to the release of EVs that promoted MuSC-mediated formation of multinucleated myotubes at an extent comparable to that observed with EVs from FAPs isolated from mdx mice treated *in vivo* with TSA (Fig. 2A and B – compare TSA vitro to untreated). These results are consistent with data from human DMD patients shown in Fig. 1A and B.

**Fig. 2.**
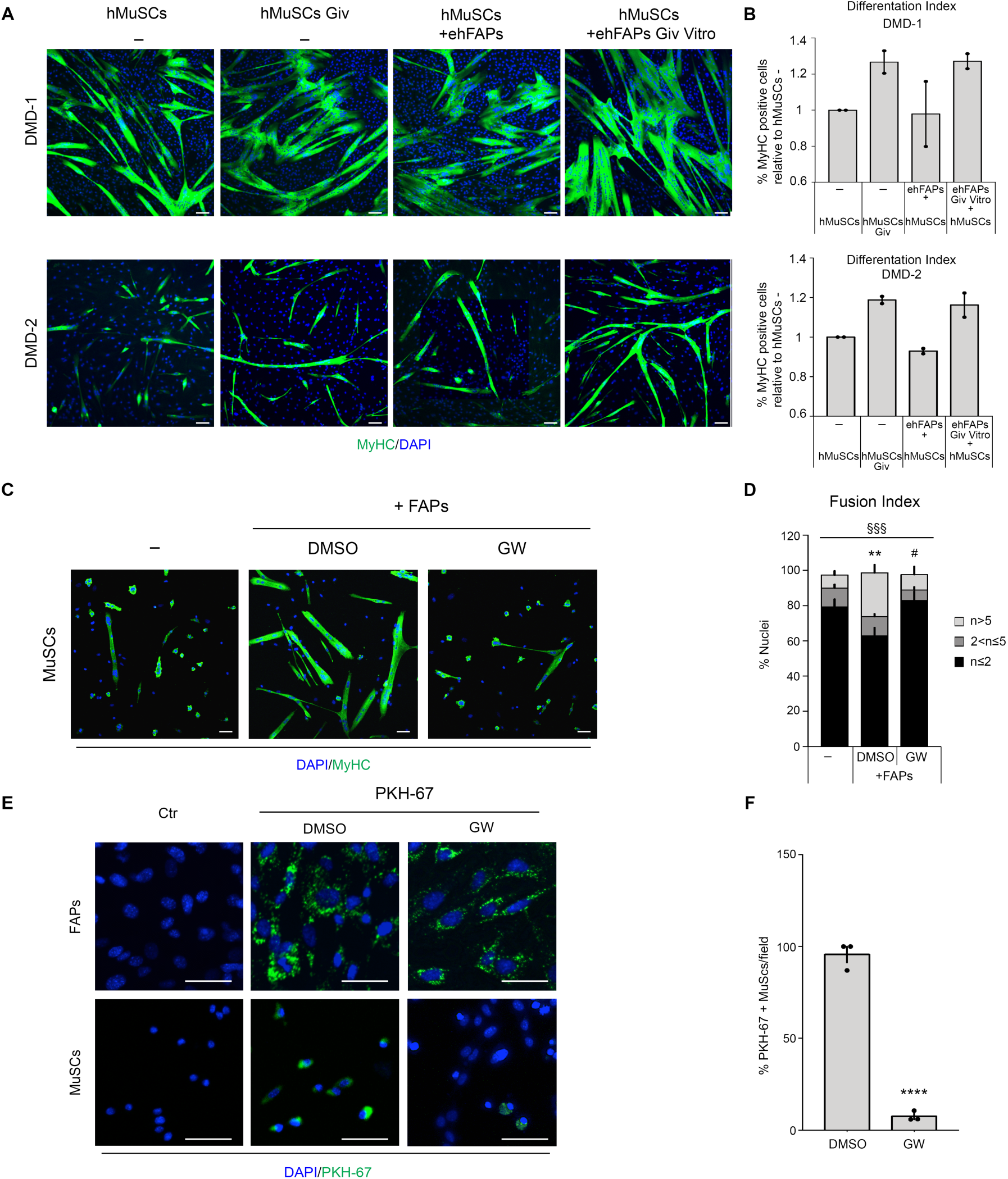
EVs from HDACi-treated FAPs promote MuSC differentiation into multinucleated myotubes. **(A)** Representative images of myogenic differentiation of mdx MuSCs, as assessed by immunostaining for MyHC (green). MuSCs were cultured either alone (-) or incubated with EVs isolated from 1.5 month old mdx mice (EVs FAPs) either treated with vehicle (CTR) or exposed to TSA *in vivo* (TSA). Alternatively, mdx FAPs were exposed *in vitro* to TSA (TSA vitro) prior to co-culture with MuSCs. Scale bar = 50 μm. **(B)** Graph showing the fusion index of MuSCs in the condition described in A, (n=3). Star (*) indicates statistical analysis by Tukey test on the % of n>5 (% of nuclei that were MyHC+ inside myotubes containing more than 5 nuclei) relative to MuSCs cultured alone (-), **p <0.01, ****p>0,0001; Hash (#) indicates statistical analysis by Tukey test relative to EVs FAPs CTR, ##p <0.01, ###p<0,001; ns= not significant; § represents statistical analysis by 2way Anova test, §§§§p < 0,0001. Nuclei were counterstained with DAPI (blue). All data correspond to the average ± SEM. **(C)** Flow cytometry analysis to count EVs isolated from mdx FAPs, cultured and treated in vitro with DMSO (CTR) or TSA (50nM for 12hrs) (TSA vitro) and stained with PKH-67 lipidic dye. The upper panels from left to right are representing unmarked control (EVs-FAPs NM), PKH-67 positive microvesicles in the supernatant of DMSO treated FAPs (EVs-FAPs CTR PKH-67) and PKH-67 positive microvesicles in the supernatant of TSA *in vitro* treated FAPs (EVs-FAPs TSA vitro PKH-67). The lower panels, from left to right, are showing the Flow Cytometer calibration beads (the numbers represent the size of the beads, expressed in nanometers, corresponding to the peak) and the dimension of positive microvesicles isolated from the supernatant of FAPs treated *in vitro* with DMSO (EVs-FAPs CTR PKH-67) or TSA (EVs-FAPs TSA vitro PKH-67) **(D)** Graph showing the absolute number of PKH-67 positive EVs of size ranging from 110 to180 nm isolated from FAPs exposed in vitro to vehicle (CTR) or TSA (n=3). Star (*) indicates statistical analysis by t-test relative to EVs-FAPs CTR; *p>0,05. For TSA in vivo treatments, mice were i.p. injected daily with 0.6 mg/kg of TSAfor 15 days. For TSA in vitro treatment: FAP cells were exposed to TSA (50nM) for 12hr.

We also evaluated whether HDACi could increase the abundance of FAP-derived EVs. To this purpose, we exposed the same number of FAPs isolated from vehicle-treated mice to TSA *ex vivo,* and quantified the number of EVs released in the supernatant. Fig. 2C and D show that TSA increased the abundance of EVs in the supernatant of FAP cultures, as compared to CTR-treated FAPs. As the supernatant was collected from the same number of cells in each condition, the increased amounts of EVs did not depend on the effect of TSA on cell proliferation, but likely reflected an increased cellular output of EVs.

### Enrichment of miR206 in interstitial EVs correlates with compensatory regeneration of DMD muscles and treatment with HDACi

We next investigated whether FAP-derived EVs could be detected in regenerating muscles, in physiological (acute injury) or pathological (muscular dystrophy) conditions. To this purpose, we performed *in situ* immunofluorescence for CD63, an integral membrane protein enriched in EVs, on muscle sections from young Wild Type (WT) unperturbed (CTR) or regenerating muscles (4 days post injury-cardiotoxin injection, CTX), and from mdx mice at different stages of disease progression (i.e. 1.5-month or 12-month old mice). We also included sections of muscles from mdx mice that were exposed or not to the TSA, which reportedly promotes regeneration in young, but not old, mdx mice(*18, 20*). This analysis revealed sporadic CD63 signal in WT unperturbed and old mdx muscles, while a dramatic increase in CD63 signal was observed in regenerating WT and mdx young muscles (fig. S2A and B). Interestingly, in all experimental conditions the vast majority of CD63 signal was invariably detected in the interstitium between regenerating, embryonal eMyHC-positive fibers, and largely overlapped with interstitial Sca1 signal, which identifies putative FAP cells. Notably, muscles of mdx young mice exposed to TSA showed a significant increase in CD63-positive interstitial signal, as compared to young mdx CTR muscles, with levels comparable to those observed in regenerating WT muscles (fig. S2A and B). By contrast, TSA treatment could not increase the CD63 signal in the interstitium of muscles from 1 year-old mdx mice, which have been previously shown to be resistant to the beneficial effects of HDACi(*18, 20*). The overlap between interstitial CD63 and Sca1 signals, the close proximity to eMyHC-positive regenerating myofibers, together with the differential response of young versus old mdx muscles to HDACi, suggest that FAP-derived EVs could be implicated in HDACi-mediated activation of MuSCs to regenerate dystrophic muscles, as previously indicated by transwell co-cultures experiments(*18*).

Previous works have implicated microRNA (miRs), as mediators of exosome-regulated biological processes(*31–36*). To investigate miRs contribution to the ability of FAP-derived EV to regulate MuSC activity, we first determined whether FAP-derived EVs could transfer RNA to MuSCs, by using fluorescent acridine orange (AO) - a specific nucleic acid staining. AO labeled FAPs-derived EVs were incubated with freshly isolated MuSCs. Confocal microscopy analysis revealed AO-labelled intracellular RNA dots inside recipient MuSCs, both within and outside the nuclear membrane (fig. S3A). We have previously shown that HDACi extensively change the miR expression pattern in FAPs from mdx muscles(*20*). Therefore, we sought to further investigate the functional impact of siRNA-mediated blockade of miR biogenesis on muscle regeneration. We evaluated the effect of siRNA-mediated knockdown of Drosha, the RNA-specific endoribonuclease required for microRNA biogenesis, on functional interactions between FAPs isolated from HDACi-treated young mdx mice and MuSCs isolated from control-treated young mdx mice. siRNA efficiently down-regulated (by about 60%) Drosha expression in FAPs (Fig. 3A), and drastically reduced the ability of TSA-treated FAPs to enhance MuSC-mediated formation of multinucleated myotubes in transwell co-culture (Fig. 3B and C). We then determined the identity of the miRs within FAP-derived EVs, by performing a Taqman-based miR expression microarray using EVs purified from the supernatant of FAPs isolated from mdx mice treated for 15 days with TSA or control vehicle (CTR). This analysis identified several miRs up-regulated in EVs from TSA-treated mdx mice. We annotated the top 14 miRs significantly up-regulated in FAPs from TSA-treated mdx mice (Fig. 3D and Fig. S3B), including miRs that have been previously implicated in the regulation of skeletal myogenesis (*35*),(*37–43*). Ingenuity Pathway Analysis (IPA) revealed a number of regulatory pathways potentially affected by these miRs and implicated in the control of MuSCs (Fig. S3C). Interestingly, RNAseq data generated from MuSCs isolated from mdx mice treated with TSA versus control vehicle revealed the activation of top pathways (*Notch*, *JAK-STAT*) that were also predicted by IPA analysis of up-regulated miR in EVs from FAPs of TSA-treated mdx mice (Fig. S3D). Among the TSA-induced EV-miRs, the most up-regulated (17.06 fold) was the muscle-specific (myomiR) miR206-3p (herein indicated as miR206) - a central component of skeletal muscle regeneration(*43, 44*), and HDACi-activated network in dystrophic muscles(*20, 45*). Of note, transgenic over-expression of miR206 has been successfully exploited as tool for the treatment of DMD and other muscular disorders(*36, 43, 46*). Interestingly, *in situ* hybridization analysis of muscle biopsies from DMD patients at various ages revealed increased levels of interstitial miR206, as compared to control biopsies (from non DMD boys) (Fig. 3E and F). This analysis also revealed a progressive reduction of interstitial miR206 along with the disease progression in coincidence with the exhaustion of the regenerative activity and the increased severity of fibrosis (Fig. 3G-J). The same pattern could also be observed in mdx mice, in which an abundant miR206 signal was detected in the interstitium of tibialis anterior muscles, overlapping with the immunofluorescence staining of the FAP’s marker Sca-1 (Fig. 4A and B).

**Fig. 3:**
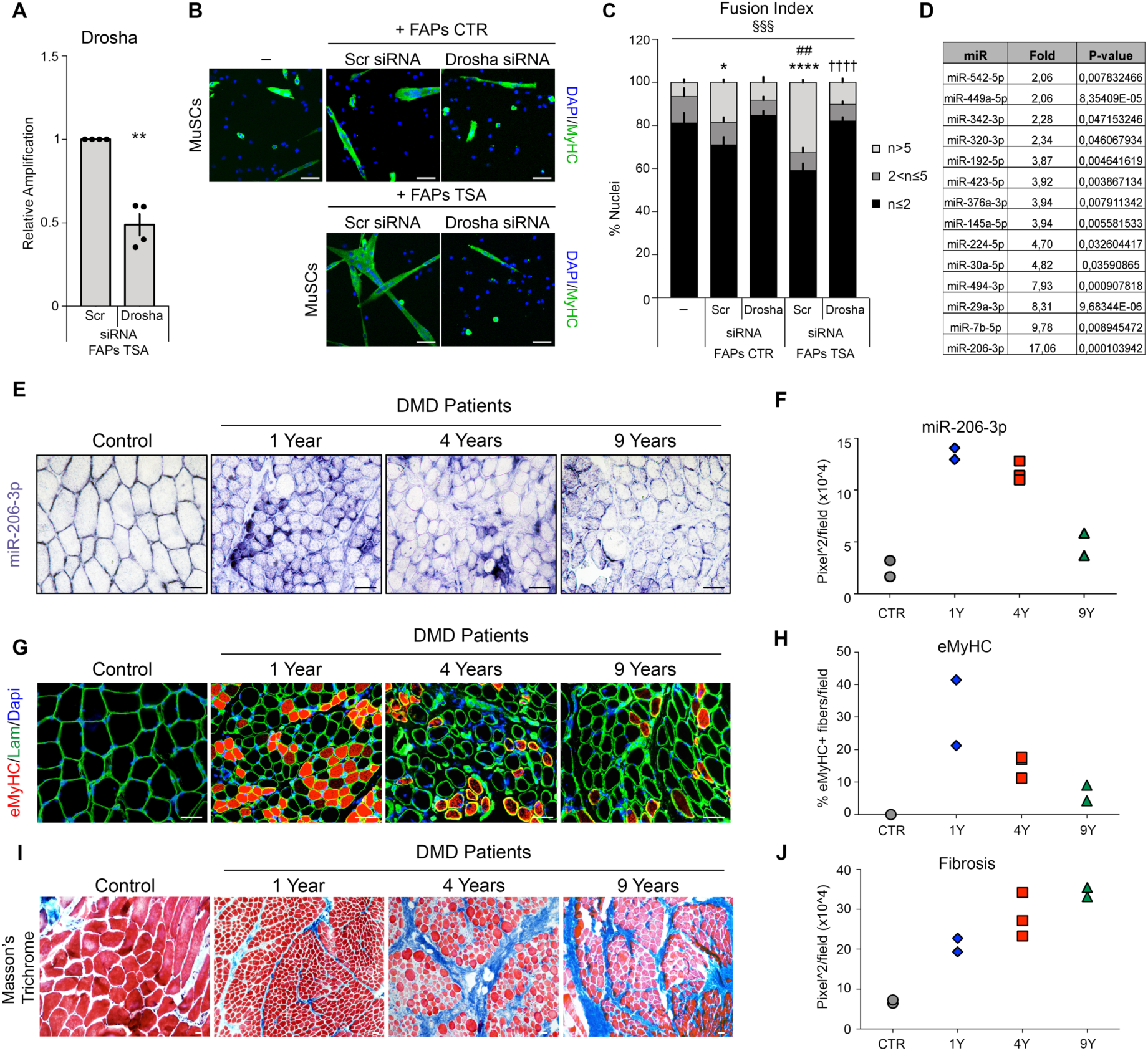
Increased amounts of miR206-3p in DMD muscles, associate with muscle regeneration and inversely correlate with disease progression. **(A)** Graph showing the relative expression of Drosha in FAPs after Drosha down-regulation by siRNA (n=4). Star (*) indicates statistical analysis by t-test relative to FAPs TSA treated and transfected with scramble (Scr); **p<0.01. **(B)** Representative images of myogenic differentiation of MuSCs assessed by immunostaining for MyHC (green). MuSCs were cultured alone (-) or in transwell co-culture with FAPs isolated from mdx mice treated either with vehicle (+FAPs CTR) or TSA (0.6 mg/kg/day for 15 days by i.p.) (+FAPs TSA), and transfected with scramble (Scr siRNA) or Drosha siRNA (Drosha siRNA) prior to co-culture with MuSCs. Scale bar = 50 μm. **(C)** Graph showing the fusion index of MuSCs in the conditions described in B (n=4). Star (*) indicates statistical analysis by Tukey test relative to MuSCs alone (-); *p<0.05, ****p<0.0001. Hash (#) indicates statistical analysis by Tukey test relative to MuSCs in co-culture with FAPs CTR transfected with scramble (Scr); ##p < 0.01. Cross (†) means Tukey analysis compared to MuSCs in co-culture with FAPs TSA transfected with scramble (Scr): ††††p < 0,0001. § represents statistical analysis by 2way Anova test. §§§p < 0,001. **(D)** Table representing a manually assembled list of microRNAs revealed by microarray analysis and statistically induced by TSA in FAPs-derived EVs. **(E)** Representative images of miR206-3p (violet) immunohistochemistry in *vastus medialis* bioptic samples of Control and DMD patients at different ages: 1 year (n=2), 4 years (n=3) and 9 years old (n=2). **(F)** Graph representing miR206-3p violet area quantification relative to E. **(G)** Representative images of immunofluorescence for eMyHC (red) and Laminin (green condition described in E. **(H)** Graph showing the percentage of eMyHC positive fibers of condition described in G. **(I)** Representative images of Masson’s Trichrome staining in condition described in E. **(J)** Graph showing the fibrotic area quantification of conditions described in I. For E, G, I Scale bar = 50 μm.

**Fig. 4:**
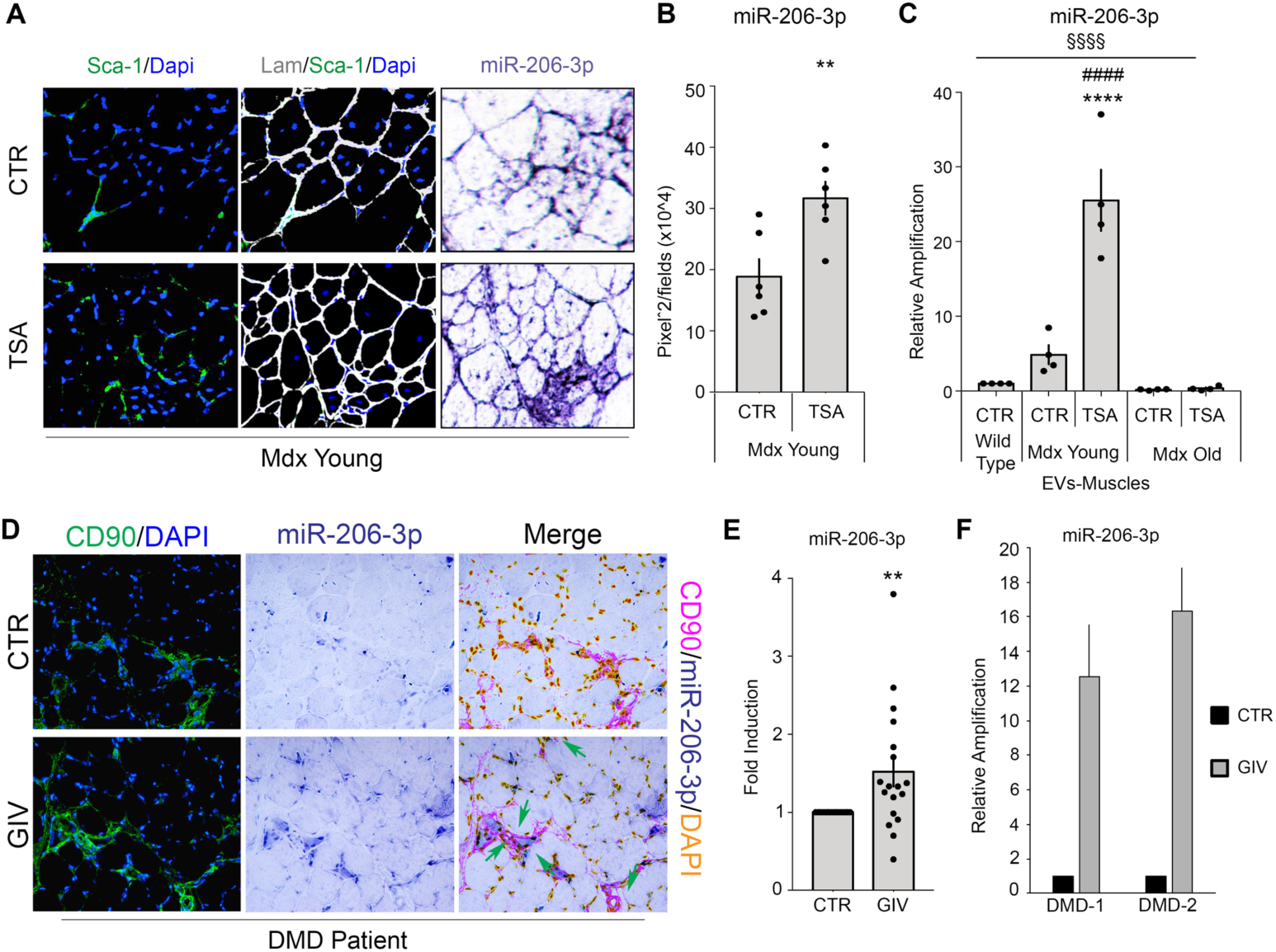
HDACi increase miR206-3p amounts in EVs from DMD FAPs. **(A)** Representative images of Sca-1 (green), Laminin (white), DAPI (blue) immunofluorescence, and miR206-3p (violet) immunohistochemistry on sequencial cryosections of Tibialis Anterior (TA) of young mdx mice (CTR) (n=6) and after TSA treatment (TSA) (0.6 mg/kg/day for 15 days by i.p.) (n=6). **(B)** Graph showing the quantifications of miR206-3p violet area relative to conditions indicated in A. Star (*) indicates t-test analysis **p<0.01. **(C)** Graph representing the miR206-3p relative expression in EVs isolated from muscle interstitium of tibialis anterior from wild type mice (CTR), mdx young and old mice (control –CTR- and TSA treated) (n=4). Star (*) indicates Tukey analysis compared to wild type mice (CTR), *p<0.05 ****p < 0.0001; hash (#) indicates Tukey analysis compared to mdx young CTR, #### p<0.0001. § represents statistical analysis by Anova test. §§§§ p<0,0001. **(D)** Representative images of CD90 (green), Dapi (blue) and miR206-3p (violet) immunohistochemistry in brachial *biceps* bioptic samples of DMD patient before (CTR) and after Givinostat treatment (GIV). **(E)** Graph showing the quantifications of miR206-3p violet area measured as pixel^2/field relative to the experimental points indicated in D (n=18). Star (*) indicates t-test analysis. **p<0.01. **(F)** Graph showing the relative expression of miR206-3p in EVs isolated from a human population of muscle resident cells enriched in FAPs (ehFAPs) isolated from biopsies of two DMD patients (DMD-1 and DMD-2) before (CTR) and after (GIV) Givinostat treatment. Nuclei were counterstained with DAPI (blue). All data for B, C and E correspond to the average ± SEM while the data for F correspond to the average ±SD.

We next investigated whether exposure to HDACi could increase the interstitial amount of intra-vesicular miR206 to the levels observed at earlier stages of disease progression. Treatment of young mdx mice (1.5 months of age) with TSA, which promotes regeneration and inhibits fibrosis (*18, 20*), could increase the amount of interstitial miR206 signal associated with Sca-1 staining, which was used to identified putative FAPs (Fig. 4A and B). We performed a quantitative assessment (by qPCR) of EV-miR206, upon isolation of whole tibialis anterior (TA) muscles from TSA-treated mdx muscle or control-treated mdx mice, followed by mechanic dissociation and EVs purification(*47*). This procedure enriches for interstitial material, as it minimizes the EVs content in myofibers, and showed a ∼5 fold increase of miR206 in TSA-treated mdx muscle as compared to control-treated (Fig. 4C).

We then performed *in situ* hybridization analysis of miR206 expression and localization in muscle biopsies from DMD patients before and after the treatment with Giv. We used muscle sections available from 18 patients, among the 19 DMD boys enrolled in this clincal trial (as one dropped out of trial). In these patients we found an increased abundance of miR206 after Giv treatment (Fig. 4D and E). Of note, the miR206 signal was predominantly detected in the interstitial space, and was almost invariably juxtaposed to CD90 signal, which identifies human FAPs (see merge picture, in which the IF and hybridization signals are merged, the CD90 signal turns purple and the miR206 signal remains violet). This suggests that FAPs are the main source of the interstitial miR206 in DMD muscles. Consistently, we detected 12 to 16 fold increase of miR206 in the EVs from ehFAPs of the two DMD biopsies available (Fig. 4F).

### EV-miRs cooperatively promote compensatory regeneration while reducing fibrosis and inflammation in dystrophic muscles

The enrichment of interstitial EVs with increased amounts of miR206 in regenerating DMD muscles suggests that FAP-derived EVs could transfer miR206 to MuSC to promote compensatory regeneration. While miR206 transfer by exosomes from MuSCs to interstitial fibroblasts has been recently described as a mechanism to hamper the formation of fibrotic scars during skeletal muscle hypertrophy(*36*), the reciprocal process – that is, transfer of miR206 from FAPs to MuSCs - has not been described so far. We therefore explored the potential transfer of miR206 from FAPs to MuSCs, through EV, within the context of the mdx mice. To this purpose we quantified MuSC’s uptake of FAP-derived EVs *in vivo*, by tracing EVs with PKH-67 labelling. EVs were first purified from FAPs isolated from mdx mice previously treated with TSA (EVs-FAPs TSA) and then exposed to PKH-67, prior to their transplantation into tibialis anterior (TA) muscles. FACS analysis showed that 17.2% of the whole population of cells isolated from transplanted TA muscle uptaked PKH-67-labelled EVs (Fig. 5A, left panel). Among them, we found that about 2% of FACS-isolated MuSCs incorporated PKH-67-labelled EVs (Fig. 5A, right panel, and Fig. 5B). We argue that such a low percentage of PKH-67-positive MuSCs detected under-estimates the actual rate of MuSC incorporation of FAP-derived EVs, as it is intrinsically biased by the loss of PKH-67-positive MuSCs once they differentiate into myofibers. Consistent with EV-mediated transfer of miR206 from FAPs to MuSCs in DMD muscles exposed to HDACi, MuSCs that incorporated PKH-67-labelled EVs exhibited a five-fold increase in miR206 (Fig. 5C), further indicating that FAP-derived EVs significantly increase the intracellular amount of miR206 in MuSCs. Furthermore, global inhibition of miR biogenesis in FAPs, by Drosha knockdown via RNA interference (RNAi) prior to their EVs injection, was sufficient to prevent the increase of miR206 in PKH-67-positive MuSCs isolated from transplanted muscles (Fig. 5C). Likewise, in a parallel experiment, inhibition of EV biogenesis by exposure to GW4869 abrogated miR206 transfer from TSA-treated mdx FAPs to co-cultured MuSCs transwell plates (fig. S4A). Considering the previous report from Fry et al.(*36*), this finding is consistent with a model of reciprocal exchange of miRs between FAPs and MuSCs.

**Fig. 5:**
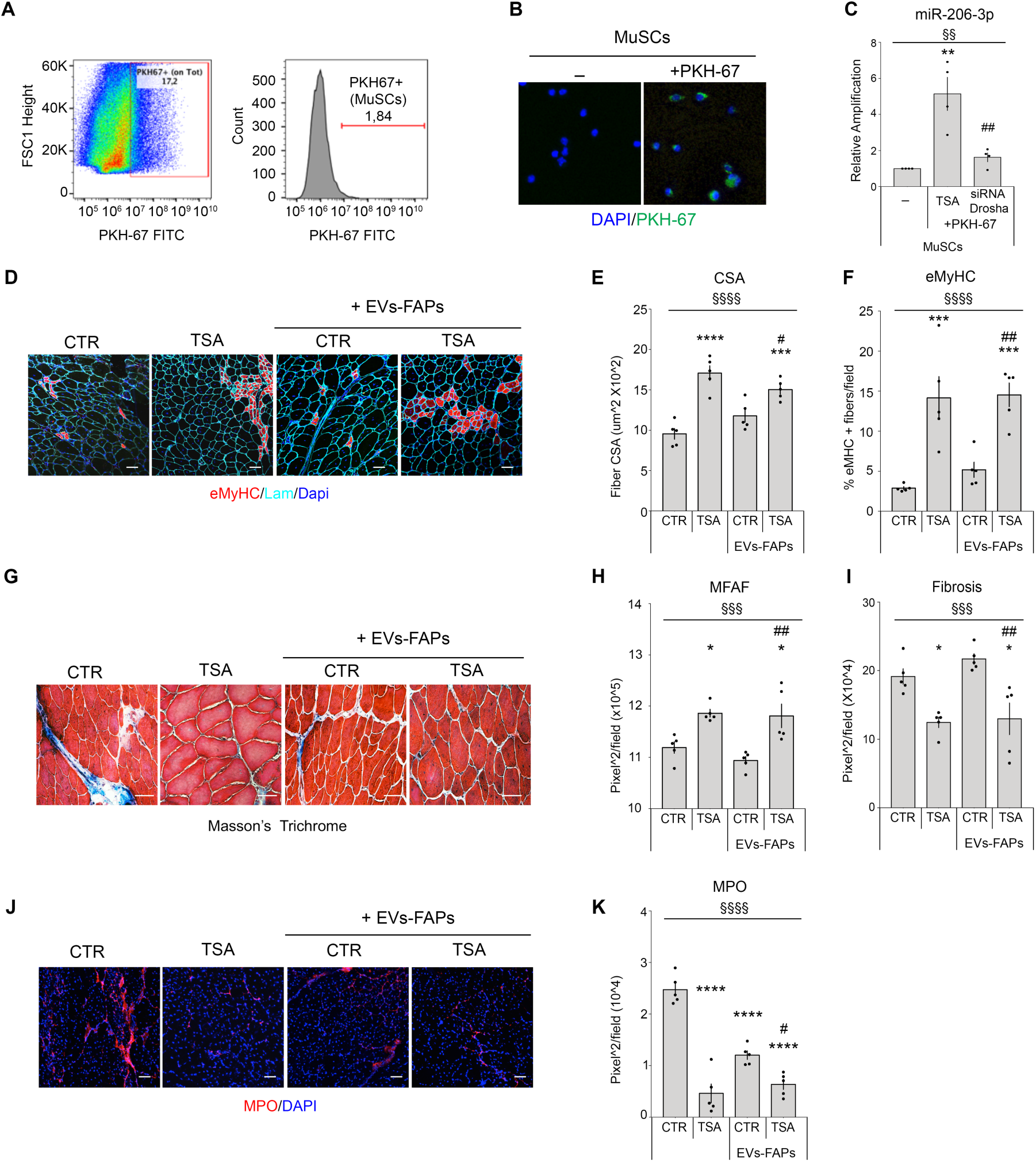
FAPs derived EVs promote regeneration and inhibit fibrosis and inflammation in dystrophic muscles of mdx mice. **(A)** Flow cytometry analysis of mdx muscles injected with PKH67-labeled EVs isolated from FAPs of 1.5 month old mdx mice exposed to TSA. The left panel shows the percentage of PKH-67 positive cells in whole muscle. The right panel shows the percentage of the PKH-67 positive MuSCs. **(B)** Representative images of PKH-67 (green) and DAPI (blue) staining in MuSCs freshly isolated from muscles previously injected with PKH67-labeled EVs. **,(C)** Graph showing the relative expression of miR206 in MuSCs isolated as described in A and B and in MuSCs from muscles injected with PKH67-labeled EVs isolated from FAPs of 1.5 month old mdx mice exposed to TSA, and then transfected or not with Drosha siRNA (siRNA Drosha) (n=4). Star (*) means significance relative to MuSCs that not uptake PKH67-labled EVs, **p < 0.01. Hash (#) means significance compared to MuSCs that uptake EVs-FAPs TSA, ^##^p < 0.01. § indicates Anova analysis, §§p<0,01. **D-K)** Stainings and relative measurements on Tibialis Anterior muscle transversal sections of 1.5 months old mdx mice treated daily for 21 days with intra-peritoneal injection of vehicle (CTR) or TSA, or once a week with intramuscular (Tibialis Anterior) transplantation of EVs derived from FAPs (EVs-FAPs) exposed or not to TSA *in vivo* (EVs-FAPs CTR, EVs-FAPs TSA). EVs were injected every seven days and sacrificed after 21 days of treatment. (n=5)**. D** Representative images of immunofluorescence for embryonic myosin heavy chain (eMyHC-red) and laminin (Lam-cyan) stainings. Scale bar = 50 μm. **E** Graph showing the quantification of cross-sectional area (CSA). **F** Graph showing the quantification of muscle regeneration (eMyHC). **G** Representative images of Masson’s Trichrome staining. Scale bar = 50 μm. **H** Graph showing the quantifications of muscle fiber area fraction (MFAF). **I** Graph showing the quantifications of fibrotic area. **(J)** Representative images of myeloperoxidase staining (MPO-red). Scale bar = 50 μm. **K** Graph showing the quantifications of mieloperoxidase (MPO). Nuclei were counterstained with DAPI (blue). Statistical analysis by Anova and Tueky test. All data correspond to the average ± SEM. In D-K Star (*) means significance compared to CTR, *p<0,05; ***p<0,001; ****p<0,0001; while hash (#) means significance compared to EVs-FAPs CTR, #p<0,05, ##p<0,01. § indicates significance by Anova test; §§§ p<0,001; §§§§ p<0,0001. TSA in vivo was administrated 0.6 mg/kg/day for 21 days by i.p.

We next investigated the functional impact of this process, by monitoring the effect of EVs-FAPs TSA transplantation on parameters of disease progression in mdx mice. Since HDACi exert beneficial effects in mdx mice by promoting compensatory regeneration and by reducing fibrosis and inflammation(*16*), we sought to evaluate to what extent these effects are accounted by FAP-derived EVs. To this purpose, we compared the ability of intramuscular (i.m.) injection of EVs purified from FAPs isolated from mdx mice that were previously treated with TSA (EVs-FAPs TSA) or vehicle (EVs-FAPs CTR) to stimulate regeneration, reduce fibrosis and inflammation, as compared to systemic exposure to TSA (or vehicle control), which was administered via daily intra-peritoneal injection for 21 days. The EVs i.m. injection was repeated every 7 days for 21 days, in order to allow a direct comparison with systemic expsore to TSA. EVs-FAPs TSA injected muscles showed an increase in embryonic-MyHC-positive myofibers (eMyHC) and cross sectional area (CSA), comparable to the increase observed in the muscle following systemic exposure to TSA (Fig. 5D-F). Likewise, EVs-FAPs TSA i.m. injection and systemic delivery of TSA showed a comparable ability to reduce fibrosis and increase muscle fiber area fraction (MFAF) (Fig. 5G-I and fig. S4B,C), as well as inflammation, as determined by reduction of myeloperoxidase (MPO)-positive areas, in mdx muscles (Fig. 5J-K).

We also investigated whether the HDACi-mediated up-regulation of miRs in FAP-derived EVs is direct and cell specific. We found that direct exposure of FAPs to TSA *ex vivo* induced an up-regulation of the same miRs detected by microarray from EVs isolated from the supernatant of FAPs derived by TSA-treated mdx mice *in vivo* (see Fig. 3D), with a comparable magnitude (fig. S4D). Moreover, *ex vivo* exposure to TSA was used to determine whether HDACi-induced up-regulation of miR206 in EVs could be observed also in muscle-derived cell types that typically express miR206 – i.e. MuSCs and myotubes. Fig. S4E-H shows that upon *ex vivo* exposure to TSA only FAP-derived EVs exhibited up-regulation of miR206, although TSA could increase the transcription of miR206 in both MuSCs and myotubes. This data indicates that transcriptional upregulation of miRs by HDACi is not sufficient to increase their levels in EVs, and points to the importance of cell type-specific post-transcriptional regulation of miR incorporation into EVs.

### EVs isolated from FAPs of HDACi-treated dystrophic muscles are required to promote MuSCs activation and amplification

To gain mechanistic insights into the pro-myogenic activity of EVs derived from FAPs of HDACi-treated mdx mice, we searched for putative target genes of miR206 that matched with downregulated genes in our RNA-seq datasets obtained from MuSCs of TSA-treated mdx mice. A downstream analysis was carried only for miR206 targets that have been already annotated or predicted with high confidence. Ingenuity pathway analysis (IPA) revealed a potential effect of miR206 on various signaling pathways involved into muscle development and disease, including *Utrophin* (UTRN), *Pax7* and *Notch3*(*48–51*) (fig. S5A). We decided to focus on miR206 regulation of *Notch3*, as negative regulator of MuSCs differentiation, because previous works demonstrated that miR206 down-regulation of *Notch3* promotes MuSC differentiation(*51*). We therefore purified EVs from FAPs isolated from TSA- or control (vehicle)-treated 1.5 month-old mdx mice for 21 days and evaluated their effect on MuSC ability to form myotubes, while monitoring the expression levels of *Notch3* and *Notch1*. EVs from FAPs of TSA-treated mdx mice increased MuSC ability to form multinucleated myotubes (Fig. 6A-B) While *Notch3* and *Notch1* transcripts were both reduced in MuSCs isolated from mdx mice treated with TSA, we observed a selective down-regulation of *Notch3* transcripts in MuSCs isolated from control-treated mice and cultured with EVs derived from FAPs of TSA-treated mdx mice (Fig. 6C and D).

**Fig. 6:**
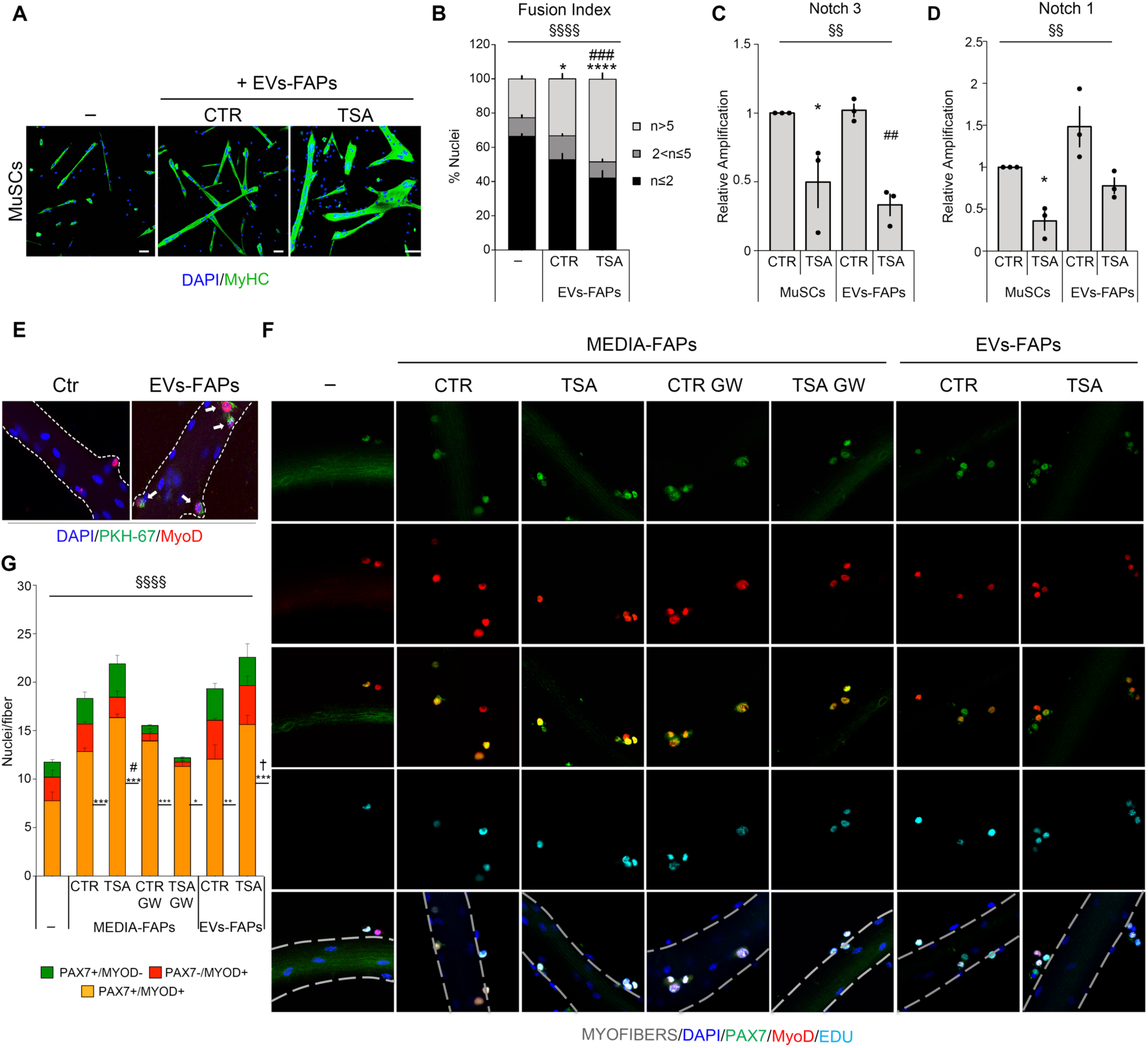
EVs from FAPs *in vivo* exposed to TSA promote MuSCs differentiation, expansion and asymmetric division. **(A)** Representative images of myogenic differentiation of MuSCs assessed by immunostaining for MyHC (green). Nuclei were counterstained with DAPI (blue). MuSCs were cultured alone (-) or incubated with EVs (EVs FAPs) isolated from FAPs of 1.5 month old mdx mice exposed to veichle (CTR) or TSA (TSA) (n=7). Scale bar = 50 μm. **(B)** Graph showing the fusion index of MuSCs in the condition described in A. Star (*) indicates statistical analysis by Tukey test relative to MuSCs cultured alone (-), *p<0.05, ****p>0,0001; hash (#) means significance compared to EVs-FAPs CTR, ^###^p<0,001. § indicates significance by 2way Anova test; §§§§ p>0,0001. **(C)** Graph relative to Notch3 expression in MuSCs isolated from 1.5 month old mdx mice treated either with vehicle (CTR) or TSA (TSA), and in EVs derived from FAPs (EVs-FAPs) isolated from 1.5 month old mdx mice treated either with vehicle (CTR) or TSA (TSA) n=3). Star (*) indicates statistical analysis by Tukey test relative to mdx vehicle treated mdx mice (CTR); *p<0,05;. hash (#) means significance compared to EVs-FAPs CTR, ^##^p<0,01. § indicates significance by Anova test; §§ p>0,01. **(D)** Graph relative to Notch1 expression in MuSCs isolated from 1.5 month old mdx mice treated either with vehicle (CTR) or TSA (TSA), and in EVs derived from FAPs (EVs-FAPs) isolated from 1.5 month old mdx mice treated either with vehicle (CTR) or TSA (TSA) n=3). Star (*) indicates statistical analysis by Tukey test relative to mdx vehicle treated mdx mice (CTR); *p<0,05;. hash (#) means significance compared to EVs-FAPs CTR; *p < 0.05. § indicates significance by Anova test; §§ p>0,01. **(E)** Representative images showing myofibers cultured alone (Ctr) or with extracellular vesicles isolated from FAPs (EVs-FAPs) and stained with PKH-67 (green) and MyoD (red). Nuclei were counterstained with DAPI (blue)**. (F)** From left to right: representative images of myofibers cultured for 24 hours alone (-) or with conditioned media of FAPs that were previously exposed or not to TSA and GW4869 (MEDIA-FAPs) or extracellular vesicles (EVs-FAPs) collected from FAPs isolated from 1.5 month old mdx mice treated either with vehicle (CTR) or TSA (TSA). Myofibers were stained for EdU (Cyan), immunofluorescence with Pax7 (green), MyoD (red). Nuclei were counterstained with DAPI (blue). (n=3). **(G)** Graph showing the average of nuclei positive for Pax7 (green), MyoD (red) in the conditions indicated in F. Star (*) indicates statistical analysis by Tukey test relative to myofibers cultured alone (-), (n=4).*<p0,05; **p<0,01 ***p < 0.001, hash (#) means significance compared to Media-FAPs CTR, ^#^p<0,05; cross (†) means significance compared to EVs-FAPs CTR, ^†^p<0.05. § means significance by 2way Anova test; §§§§ p < 0.0001. All data correspond to the average ± SEM. TSA in vivo was administrated 0.6 mg/kg/day for 15 days by i.p.

In keeping with the opposite roles of Notch members in MuSCs activation(*52*), we reasoned that selective down-regulation of *Notch3* by FAP-derived vesicular-miR206 could promote MuSC activation and self-renewal. To investigate this hypothesis, we evaluated the effect of EVs derived from FAPs of HDACi-treated or control-treated mdx mice on MuSCs within single fibers isolated from WT mice. Freshly isolated single muscle fibers from tibialis anterior, extensor digitorum longus, gastrocnemius and soleus of C57BL6J mice were incubated with FAP-derived EVs or whole conditioned media, and their effect on MuSCs was evaluated by monitoring the expression of Pax7, MyoD and DNA synthesis (EdU incorporation). In this experimental setting, the uptake of FAP-derived EVs by MuSCs from single fibers was first confirmed by staining of lipidic dye PKH-67 used to label purified vesicles (Fig. 6E). Fig. 6F and G show that the whole supernatant derived from FAPs of control-treated mdx mice increased the number of Pax7+/MyoD+ MuSCs that exhibited enhanced DNA synthesis (Edu+), and could be therefore considered activated MuSCs(*53*). Indeed, nearly 100% of the EdU+ nuclei were also Pax7+/MyoD+. Moreover, the whole supernatant derived from FAPs of TSA-treated mdx mice could further augment this effect. While a drastic reduction in the number of quiescent (Pax7+ MyoD-) and differentiation-committed MuSCs (Pax7-MyoD+) was observed upon incubation with the whole supernatant of FAPs isolated from control-treated mdx mice and pre-exposed to GW4869, the number of Pax7+/MyoD+ MuSCs with high DNA synthesis (Edu+) was not altered in these conditions (Fig. 6F and G). By contrast, GW4869 treatment of FAPs from TSA-treated mdx mice drastically reduced the ability of the whole supernatant to increase all MuSC populations as well as their proliferation (Fig. 6F and G). This data indicates that the ability of FAPs from HDACi-treated mdx mice to promote expansion of MuSCs depends on EVs. Indeed, only EVs from FAPs of TSA-treated (but not from control-treated) mdx mice increased the number of Pax7+/MyoD+ MuSCs (Fig. 6F and G)

## Discussion

Defining the identity of extracellular mediators that coordinate the activities of the cellular components of the muscle stem cell niche is a major challenge in regenerative medicine, as it might inspire selective interventions toward promoting compensatory regeneration of diseased muscles, while eliminating the most unfavorable outcome of disease progression – the maladaptive repair by deposition of fibrotic and adipose tissue that typically compromises contractile activity and regenerative potential of muscles.

While a variety of soluble factors, including inflammatory cytokines and growth factors, have been implicated as mediators of reciprocal functional interactions between the major cellular components of the muscle stem cell niche, EVs are emerging as powerful tool to transfer biochemical and genomic information from one cell type to another(*54, 55*). Indeed, by virtue of their content in proteins and RNA, EVs enable the cross-activation of biological process between neighbor cells that accounts for their functional interactions. As such, the qualitative and quantitative control of proteins and RNA content appears as two major determinants of EVs biological activity. Likewise, the direction of cell-to-cell transmission of EVs influences the final biological outcome of EVs activity. Our finding that a defined collection of miRs in EVs derived from FAPs of dystrophic patients and mice exposed to HDACi promote MuSC activity and compensatory regeneration, while inhibiting fibrosis and inflammation in mdx mice, begins to elucidate fundamental aspects of EVs biology within the context of FAP-MuSC interactions during muscle regeneration and DMD.

A seminal discovery from Peterson’s lab has recently shown that MuSC-derived exosomes containing miR206 are required to eliminate fibrosis during muscle hypertrophy(*36*). Moreover, myofiber-derived exosomes have been reported to promote skeletal myogenesis(*56*). The pro-regenerative and anti-fibrotic activities of miR206 have been largely documented by previous studies(*44*). However, while these studies invariably indicated skeletal myofibers and MuSCs as only source of miR206, our previous studies have revealed that FAPs from dystrophic muscles turn into an alternative cellular source of miR206, once exposed to HDACi, and identified a miR206-directed network that represses the adipogenic and fibrotic program in FAPs of dystrophic muscles(*20*). The results reported in the current study further extend our knowledge on the role of FAP-derived miR206 in DMD muscle exposed to HDACi, by showing that it is incorporated into EVs, which are released in the regenerative environment. This evidence indicates the pharmacological potential of HDACi-mediated modulatation of EV content as therapeutic tool for DMD. At the same time, it raises important questions related to the function of miR206 in the EVs released by dystrophic FAPs upon exposure to HDACi, as FAP-derived EVs enriched in miR206 are paradoxically transferred to one of the major cellular sources of miR206 – the MuSCs. This raises the issue of why an external source of miR206 is required to support MuSCs activity. In this regard, it should be noted that FAP-derived EVs can deliver to MuSCs a cargo of other miRs and proteins; therefore, miR206 effects on MuSCs *ex vivo* and on regeneration of dystrophic muscles *in vivo* has to be interpreted as a combinatorial effect of the whole EV content in miRs and proteins that results in the MuSC activation and regeneration of new fibers. Notwithstanding, mdx MuSCs that incorporated EVs released from HDACi-treated FAPs exhibited a five-fold increase in miR206. In this regard, while the levels of miR206 are significantly lower in FAPs and FAP-EVs, as compared to MuSCs, repeated uptake of large amounts of FAP-derived EVs overtime can lead to an overall increase of miR levels, as observed in MuSCs from DMD muscles exposed to HDACi.

We propose a model whereby in the context of DMD additional amounts of miR206 levels from FAP-derived EVs could correct the dysfunctional phenotype of MuSCs by normalizing intracellular miR206, as also indicated by previous studies showing that transgenic expression of miR206 has beneficial effects in mdx mice(*43*). The recent discovery that dystrophin is expressed in MuSCs, and dystrophin-deficient MuSCs lose the ability to divide asymmetrically and regenerate DMD muscles(*57, 58*), indicate that additional events downstream of dystrophin can be dysregulated in DMD MuSCs. Among them, repression of miR206 expression (*45, 59*) could be compensated by FAP-derived EV miR206 and therefore restore MuSC ability to regenerate dystrophic muscles. Overall, our study provides the first evidence that FAP-derived EVs mediate functional interactions with MuSCs, and that EV-miR content could be pharmacologically modified by epigenetic drugs (e.g. by HDACi) for therapeutic purposes in both human DMD patients and mouse model of DMD. In this regard, we note that the beneficial effects of focal delivery of EVs-FAPs TSA by i.m. injection were comparable or even greater than those observed upon systemic delivery of TSA (see Fig. 5).

Further studies should elucidate the mechanism by which HDACi regulate EV biogenesis and content, and will define the potential for further development of interventions based on FAP-derived extracellular vesicles as potential selective treatment for DMD. Our study also suggests that specific interstitial amount of intra-EV miRs (i.e. miR206) can be used as biomarker of disease progression and of therapeutic response to interventions that counter DMD progression.

## MATERIALS AND METHODS

### Experimental model and subject details

#### Animals

Mice were bred, handled and maintained according to the standard animal facility procedures and all experimental protocols were approved by the internal Animal Research Ethical Committee according to the Italian Ministry of Health and the ethic committee of the Fondazione Santa Lucia (FSL) approved protocols.

C57/BL6 mice were provided by the Core Structure of the EMMA (European *Mouse* Mutant Archive, Monterotondo, Rome), C57/Bl6 mdx mice were purchased from Jackson Laboratories. Note that the term of “young” is refereed to C57BL6/mdx mice 6/8 weeks old, age which includes the temporal frame in which the beneficial effects of HDACi are restricted (*18, 20*) while the term of “old” is refereed C57BL6/mdx mice 1 year old.

#### Human samples

Human biopsies *were* collected with informed patients’ and *parental consent* in compliance with Good Clinical Practice and the Declaration of Helsinki and provided by Dr. Bertini and Dr. D’Amico (Ospedale Bambino Gesù, Rome). We used 10 μm transversal sections isolated for research purposes from vastus medialis of DMD boys, at different ages: 1 year old (n=2), 4 years old (n=3), 9 years old (n=2) and we compared with a vastus medialis section of control (no DMD) samples.

Human bioptic samples obtained from brachial biceps from DMD boys participating to the clinical trial with Giv (www.Clinicaltrials.gov; EudraCT Number: 2012-002566-12; Sponsor Protocol Number: DSC/11/2357/43) were provided by Italfarmaco S.p.A. (Milan) (23). The phase II study was approved by the local Ethics Committees and authorized by the Competent Authority of Italy. Parents of the participants provided informed written consent and each subject provided written assent before participation.

### Mice and animal procedures

Animals were used at the specified age and treated with daily intra-peritoneal injections of Trichostatin A, TSA (0.6 mg/kg/day for 15 days to obtain FAPs for co-colture and for EVs isolation or 21 days for the transplant experiments; #T8552, Sigma), dissolved in DMSO solution or in DMSO alone as vehicle control (CTR). Muscle injury was performed by intramuscular injection in Tibliais Anterior (TA) of Cardiotoxin -CTX- (10 μM, 10 mg/mL) (#L8102, Lotaxan Valence, France, http://www.latoxan.com), 4 days before mice sacrifice.

FAPs-derived EVs at a final concentration of 10 μg in 20 μL of PBS1x (0.5ug/μl) (#14190-1444, Gibco by life technologies) and its vehicle of control (20 µl PBS1x) were injected 3 times in left TA of young C57BL6/mdx mice: every 7 days for 21 days.

### Histology and in situ hybridization

The Tibialis anterior muscles were snap frozen in liquid nitrogen-cooled isopentane and then cut transversally with a thickness of 7 µm.

For Masson’s Trichrome staining to analyze fibrotic tissue, muscle cryo-sections were fixed for 20’ at 56°C in Bouin’s Solution (#HT10132, Sigma) and then stained in Working Weigert’s Iron Hematoxilin solution for 5 min (#HT1079, Sigma), washed in running tap water for 5 min and stained in Biebrich Scarlet-Acid Fucsin for 5 min (#HT151, Sigma). Sections were rinsed in de-ionized water and re-fixed in freshly made Phosphomolybdic/Phosphotungstic/dH_2_O (1:1:2) acid solution for 8 min (#HT153, #HT152, Sigma), and then they were stained in Aniline Blue solution for 5 min (#HT154, Sigma) and in acid acetic 1% for 2 min (#27221, Sigma). The slides were dehydratated in ethanol (#02860, Sigma) and xilene (#X1040, Sigma) and mounted with EUKITT (#03989, Sigma), then visualized using a Nikon Eclipse 90i; collagen fibers are stained in blue, the nuclei stained in black and the muscle tissue is stained in red. MiRNA *in situ* hybridization was performed in formaldehyde and carbodiimide (EDC)-fixed TA cryo-sections (0,16M 90 min at RT, #25952-53-8, Merck KGaA). After washes with 0,2% glycine (#G8898, Sigma) and TBS, cryo-sections were acetylated using 0,1 M triethanolamine and 0,25% acetic anhydride for 25 min at RT (respectively #90275, #A6404, Sigma). This steps are followed by a pre-hybridization using 2X of SSC, 25% formamide (#F9037, Sigma); 0,2% Triton (#X100, Sigma) 30 min at RT and by the over night hybridization at 4°C with the hsa-miR206 probe (10 pmol, #18100-01, Exiqon) dissolved in a solution of 50% formamide, 250μg/ml tRNA (#R1753, Sigma), 200μg/ml SSDNA (#D7656, Sigma), 10% dextran sulphate (#D8906, Sigma) and 2X SSC. The hybridization was followed by specific washes with SSC to eliminate non specific binding of probe (5X SSC 5min at RT, 1X SSC 15 min at 45°C, 2%BSA in 0,2X SSC 15min at 4°C, 2X SSC 5min at RT, TNbuffer 10 min at RT and TNT buffer 15’ at RT) and by the incubation of cryo-section using anti-digoxigenin-ap fab fragments (1/100, #11093274910, Roche) dissolved in TNbuffer for 2 hours at RT. To reveal the miRNA probe specific binding cryo-sections, covered from light, were incubating over night at 4°C with 0,375 mg/ml of NBT and 0,188 mg/ml BCIP dissolved in a solution of TMNbuffer (respectively #11383213001 and #11383221001, Roche).

TNbuffer is composed of 0,1 M Tris-HCL (#T1503, Sigma) and 0,15 M Nacl (#S3014, Sigma) at pH 7,5; TNTbuffer is TNbuffer with 0,1% Tween (#P1379, Sigma) while TMNbuffer is composed of 0,1 M Tris-HCL, 0,005M MgCl_2_ (#M8366, Sigma), 0,5 M NaCl and 2mM of Levamisole (#L9756, Sigma).

### Isolation of FAPs and Satellite cells (MuSCs)

hMuSCs were isolated from human biopsies as CD31 ^neg^ /CD45 ^neg^ /CD11b ^neg^ (Lin-)/CD56 ^pos^ cells; ehFAPs as Lin-/CD56 ^neg^ cells. Live/Dead staining was used to check cell vitality.

FAP cells were isolated from mdx mice as TER119^neg^/CD45^neg^/CD31^neg^/alpha7INTEGRIN^neg^/SCA-1^pos^ cells, MuSCs were isolated from mdx mice as TER119^neg^/CD45^neg^/CD31^neg^/alpha7INTEGRIN^pos^/SCA-1^neg^ cells (*18*).

Briefly, human biopsies or hind limb muscles for each mouse were minced and put into a 15 mL tube containing 4 mL of HBSS (#24020-091, GIBCO) BSA (0.2%, #A7030, Sigma) and 10 Units/ml Penicillin and 10 μg/ml streptomycin (P/S), 2 mg/ml Collagenase A (#10103586001, Roche), 2.4U/ml Dispase II (#04942078001, Roche), DNaseI 10 mg/ml (#11284932001, Roche) at 37^°^C under gentle agitation for 1hr and 30 min. The supernatants were filtered through a 100um, 70um and 40um cell strainers (#08-771-19, #08-771-2, #08-771-1, BD Falcon). Cells were spun for 15 min at 300 g at 4^°^C, the pellets were re-suspended in 0.5 mL of HBSS 1x containing DNase I and incubated with antibodies on ice for 30 min. The following antibodies were used: CD45-eFluor 450 (1/50, #48-0451-82, Leukocyte Common Antigen, Ly-5, eBiosciences), CD31-eFluor 450 (1/50, PECAM-1, #48-0311-82, eBioscience), TER-119-eFluor 450 (1/50, clone TER-119, #48-5921-82, eBiosciences), Sca1-FITC (1/50, Ly-6A/E FITC, clone D7, #11-5981-82, eBioscience), Itga7-649 (1/500, AbLab #67-0010-01), anti human CD31-FITC (#21270313, Immunotools), anti human CD45-FITC (#130-080-202, Miltenyl biotec), anti human CD11b-FITC (#130-081-201, Miltenyl biotec), anti human CD56-PE (#12-0567-42, eBioscience). HBSS was added and cells were spun for 5 min at 300 g at 4^°^C to stop the reaction.The cells were re-suspended in HBSS containing 1% DNaseI and were isolated based on size, granulosity and fluorophores levels using a FACS MoFlo HS Cell Sorter Dako Cytomation (BD) and analyzed using FlowJo.

### Cell culture

MuSCs and FAPs, either from human or mouse samples, were cultured after sorting directly in culture media: for MuSCs: 20% FBS (#16000044, GIBCO), 10% HS (#26050-070, GIBCO), 1% Penicillin-Streptomycin (#15140, GIBCO), 1% Chicken Embryo Extract (CEE, #CE-650-F, Seralab) in DMEM + Pyruvate (#41966, GIBCO) and for FAPs: BIOAMF-2 (Biological Industries). MuSCs were plated at low density on regular cell culture dishes coated with Gelatin 0.1% (#07903, Stemcell).

FAP cells were cultured in Bioamf-2 at high density (200.000 cells in 10 cm-well plate) for extracellular vesicles isolation, at low density: 40000 or 10000 cells respectively in 6 or 24 well dishes for co-culture experiments.

For HDACi *in vitro* treatment, cells were exposed to TSA (50nM) for 12hr before EVs isolation. To isolate EVs, cells reached 80-90% of confluence then, the medium was replaced with serum free DMEM (+ pyruvate + 4.5 g/l glucose + glutamate) for 24 hr.

#### Transwell co-culture experiments

MuSCs and FAP cells, both from human or mouse samples, were co-cultured by using inserts with 1.0 μm porous membrane (#353102, #353104, Falcon) to avoid direct contact between populations. Freshly sorted MuSCs were plated in the bottom of the plate, while FAP cells were plated on the upper insert.

#### Single fibers isolation and culture

Single fibers were isolated from tibialis anterior, extensor digitorum longus, gastrocnemius and soleus muscles of C57BL6J-WT mice (*60*) and cultured in proliferating medium (GM1: DMEM + pyruvate +4.5 g/l glucose + glutamate, 10% horse serum (HS), 0.5% Chicken Embryo Extract) for 24 hours, then exposed for the next 24 hrs to GM1 conditioned media derived from culture of FAPs (MEDIA-FAPs) isolated from young mdx mouse treated for 15 days with vehicle (CTR) or mdx mice treated with TSA (TSA) or media isolated from FAPs CTR and TSA pre-treated with the GW4869 to inhibit exosome biogenesis. In a parallel, independent experiment, single fibers were exposed to EVs (EVs-FAPs) purified from MEDIA-FAPs and added to GM1 for 24 hours.

#### Cell treatments

The transfection of FAPs with pCT-CD63-GFP plasmid (#CYTO120-PA-1, System Biosciences) and with a GFP-control vector (mock) in cell culture inserts was accomplished using Lipofectamine 2000 (#12566014, Thermofisher Scientific) 6 hours before MuSCs co-culture. Transwell co-cultures were maintained in GM2 medium for 24hrs and then harvested for GFP-analyses.

To decrease FAP-EVs release, GW4869 (10 uM, #D1692, Sigma) was added to FAPs culture 30’ before the co-culture setting with MuSCs. To wash away any residual trace of GW4869, FAPs media was refreshed right before co-culture.

For siRNA transfection, FAPs isolated from mdx mice, were transfected with siRNA for Drosha, using Dharmafect3.0 (#T2003-03, Thermo Fisher Scientific) according to manufacturer instructions; after 6 hours the medium was replaced with DMEM serum free for 24 hours before EVs isolation and purification.

To stain FAPs derived EVs, FAP cells were incubated with the lipidic dye PKH-67 (#P7333, Sigma) according to manufacturer instructions prior to transwell co-culture with MuSCs.

In *ex vivo* experiments FAP-derived EVs (10μg) isolated by TEIR were put in culture with MuSCs or with myofibers in the cell culture media.

For the *in vitro* treatment with TSA, cells were exposed to TSA (50nM) for 12hr.

### Immunofluorescence

For immunofluorescence analysis, cryo-sections and cells were fixed in 4% PFA for 10 min and permeabilized with 100% cold acetone (#32201, Sigma) for 6 min at −20°C or 100% cold Methanol (#32213, Sigma) for 6 min at −20° or with 0,25% Triton for 15 min at RT. Muscle sections were blocked for 1h with a solution containing 4% BSA (#A7030, Sigma) in PBS. The primary antibodies incubation were performed O.N. at 4°C and then the antibody binding specificity was revealed using secondary antibodies coupled to Alexa Fluor 488, 594, or 647 (Invitrogen). Sections were incubated with DAPI in PBS for 5 minutes for nuclear staining, washed in PBS, and mounted with glycerol 3:1 in PBS. The primary antibodies used for immunofluorescences are: rabbit anti-Laminin (1/400, #L9393, Sigma); rat anti-SCA1 (1/100, #11-5981-82 Ly-6A/E FITC, eBioscience,); mouse anti-eMyHC (1/20, #F1.652, Developmental Studies Hybridoma Bank, DSHB, http://dshb.biology.uiowa.edu/F1-652); mouse anti-MF20 (1:20, Developmental Studies Hybridoma Bank, DSHB, http://dshb.biology.uiowa.edu/MF-20), mouse anti-PAX7 (1/10, Developmental Studies Hybridoma Bank, DSHB, http://dshb.biology.uiowa.edu/PAX7), rabbit anti-MyoD-318 (#SC760, Santa Cruz Biotechnology), EDU (#C10350, Invitrogen), rat anti-CD63 PE (#143904, clone NUG-2, Biolegend) and Myeloperoxidase/MPO (1/100 #MAB3174, R&D), Anti-Human C90 (Thy-1) (1:150 #14090980 eBioscience).

#### Cytohistochemistry

To stain lipids in ehFAPS, cells were fixed in 4% PFA for 10 min and then washed with 60% isopropanol, stained with Oil red O in 60% isopropanol and rinsed with water (*18*).

### RNA preparation and RT-qPCR

Total RNA from cell cultures and from plasma was extracted with Trizol reagent (#T9424, Sigma) and 0.5–1 μg was retrotranscribed using the TaqMan reverse transcription kit (Applied Biosystems). Real-time qPCR was performed using primers (MmDrosha Fw:5’ TGCAAGGCAATACGTGTCATAG 3’; MmDrosha Rev:5’ TGAAAGCTGGTGCTGAAGGT 3’; MmNotch1 Fw: 5’TGAGACTGCCAAAGTGTTGC 3’; MmNotch1 Rev: 5’ GTGGGAGACAGAGTGGGTGT 3’; MmNotch3 Fw:5’ GTCCAGAGGCCAAGAGACTG 3’; MmNotch3 Rev:5’ CAGAAGGAGGCCAGCATAAG 3’; MmCol1a1 Fw:5’ CCTCAGGGTATTGCTGGACA 3’; MmCol1a1 Rev: 5’ GAAGGACCTTGTTTGCCAGG 3’; MmCol3a1 Fw:5’ CCCAACCCAGAGATCCCATT 3’; MmCol3a1 Rev:5’ GGTCACCATTTCTCCCAGGA 3’; MmFibronectin Fw:5’ TCCACGCCATTCCTGCGCC 3’ MmFibronectin Rev: 5’ GCACCCGGTAGCCAGTGAG 3’).

### EVs

#### EVs Isolation

EVs were isolated from FAPs serum free cell culture medium in parallel by ultracentrifugation (UC) and with Total exosomes isolation reagent -TEIR- (#4478359, Invitrogen by Thermo Fisher Scientific) according to manufacturer instructions. EVs were isolated by UC and quantified according to a previously published method(26, *27*). This isolation method included a penultimate centrifugation step in Eppendorf polypropylene conical tubes (10,000 x g for 30 min at 4°C, in Eppendorf rotor F-34-6-38) that allowed the removal/isolation of larger microvesicles. Subsequently, nano-sized EVs, comprised mainly of exosomes, were pelleted in Beckman Coulter polypropylene open top tubes (110,000 x g for 70 min at 4°C, in Beckman rotor SW28). After washing, the pellet was resuspended either in RIPA buffer or in PBS, for further immunoblotting or biophysics and molecular analyses, respectively. To estimate the amount of secreted vesicles, we quantified and compared the total protein content of the vesicle lysates using the BCA assay.

EVs were purified from tibialis anterior gently dissociated in serum-free DMEM (24 h). Cell debris and organelles were eliminated by centrifugation at 2,000g for 20 min and EVs were isolated using TEIR. For the EVs characterization by electron microscopy, the EVs were isolated using TEIR and then purified using exosomes spin columns (MW3000, Invitrogen) following the manufacturer protocol.

#### EVs characterization

EVs size distribution was determined by Dynamic Light Scattering (DLS) measurements. Collected EVs samples were diluted to a final concentration of 15μg/ml total protein content in order to avoid multiple scattering artifacts. Static and dynamic light scattering measurements were performed at 20^°^C by using a Brookhaven Instruments BI-9000 correlator and a solid-state laser tuned at *λ_0_* = 532 nm. Scattered intensity autocorrelation functions *g2(t)* have been analyzed by using multiple gamma functions for the diffusion coefficient and therefore by using the classic Stokes-Einstein relation to determine the size distribution P(D) of vesicles, where the size parameter D is actually the hydrodynamic radius of diffusing vesicles (*61*).

EVs morphology was examined for scanning electron microscopy analysis. EVs isolated and purified from FAPs by TEIR were fixed in 4% (v/v) paraformaldehyde, dehydrated by a series of incubations in 30%, 50%, and 70% (v/v) ethanol and dried on aluminium support for SEM. EVs isolated were coated with gold. A SEM LEO 1450VP (Carl Zeiss Meditec, Oberkochen, Germany) was employed to acquire backscattered electron images using 20 keV electrons leading to an information depth of about 1.5 μm. Images with a scan size of 30 × 30 μm were acquired, at a resolution of 1024 × 1024 pixels.

#### EVs labeling

##### Acridine Orange

To detect nucleic acid content, FAPs derived EVs (10μg) isolated by TEIR, were labeled with acridine orange -AO- (#235474, Sigma) (100 mg/ml) for 30 minutes at room temperature and added to MuSCs in order to reveal by immunofluorescence the RNA exchanged from FAPs to MuSCs.

##### PKH-67

To visualize EVs, FAPs-derived EVs (10μg) are stained for 5 minutes at room temperature RT with 0.5μl of PKH67 Green Fluorescent Cell Linker Kit for General Cell Membrane Labeling (#P7333, Sigma) and then polished with the exosome spin columns (#4484449, Invitrogen by Thermo Fisher Scientific) following the manufacturer protocol. The stained EVs are detected by immunofluorescence into myofibers and by cyto-fluorimetric analysis in mice muscle after their *in vivo* injection.

#### Flow cytometry analysis of extracellular vesicles released by FAPs

EVs purified from FAPs isolated from mdx mice and treated with DMSO (CTR) and TSA (50 nM), were stained with PKH67 dye and acquired on the CytoFLEX flow cytometer (Beckman Coulter, Brea, CA, USA). The cytometer was calibrated using a mixture of non-fluorescent silica beads and fluorescent (green) latex beads (Apogee, UK) with sizes ranging from 110 nm to 1300 nm. This calibration step enabled the determination of the sensitivity and resolution of the flow cytometer (fluorescent latex beads) and the size of extracellular vesicles (silica beads) (*62*) All samples were acquired at low flow rate for the same amount of time (1 min) in order to obtain an estimate of absolute counts of microvescicles comparable between various samples. The analysis of the data was performed with FlowJo software (FlowJo, LLC; Ashland, Oregon, USA). The FAPs were purified by FACS and seeded under same condition and in same number for all the samples analyzed. For EV isolation and for labeling all the samples were treated in the same way

#### EVs content

##### Protein

EVs isolated were lysed for protein extraction in RIPA buffer (50 mM Tris-HCl, pH 7.4; 150 mM NaCl; 1% NP-40; protease inhibitors). The total extravesicular protein content was quantified using the micro bicinchonic acid protein assay (BCA) (#23235, Thermo Fisher scientific).

##### Western blot

Western blot was performed on 12 ug of total lysate (WCL) and of EVs proteins using antibodies against the following proteins: rabbit anti-CD63 (1/300, H-193, Santa Cruz Biotechnology), mouse anti-Hsp70 (1/2500, clone BRM-22, Sigma-Aldrich), rabbit anti-Calnexin –CANX-(1/1000, NB100-1965, Novus Biologicals), rabbit anti-Flotillin-1 (1/200, H-104, Santa Cruz Biotechnology), mouse anti-Alix (3A9) (1/500, Cell Signaling Technologies #2171), mouse monoclonal Col3a1 (B-10) (1/150, Santa Cruz Biotechnologies #sc-271249)

As total lysate normalization we used Ponceau quantification.

##### microRNA

Total extravesicular RNA was extracted with Total Exosome RNA Protein Isolation Kit (#4478545, Thermo Fisher Scientific). TaqMan MicroRNA Assays were performed according to the manufacturer’s recommended protocols (Applied Biosystems) The threshold cycle (Ct) were defined as the fractional cycle number at which the fluorescence passes the fixed threshold. U6 snRNA served as an endogenous control for normalization.

In particular, total EVs-RNA was retro-transcribed using TaqMan MicroRNA Reverse Transcription Kit (#4366596, ThermoFisher Scientific)

For greater sensitivity, the cDNA was pre-amplified using Taqman PREAMP master mix (#4391128, ThermoFisher Scientific) and Megaplex PreAMP Primers and then amplified using high multiplexed Megaplex Primer Pool (#4444766, ThermoFisher Scientific) and the TaqMan 2X Universal Master Mix II (#4440040, ThermoFisher Scientific) on TaqMan rodent MicroRNA A/B cards Array version 3.0 (#4444909, ThermoFisher Scientific).

##### qRT–PCR

For microRNA validation, Total EVs-RNA was extracted with Total Exosome RNA Protein Isolation Kit (#4478545, Invitrogen by Thermo Fisher Scientific) and was retro-transcribed using the Qiagen reverse transcription kit (miScript II RT Kit, #218161, Qiagen) and pre-amplified using miScript PreAMP PCR Kit (# 331451, Qiagen).

Real-time qPCR was performed using (miScript SYBR Green PCR Kit, #218073, Qiagen) and using primers reported in table. All the conditions are provided in the manufacturer protocol.

### FAPs derived EVs intra-muscular injection

FAPs-derived EVs at a final concentration of 10μg in 20μL of PBS1x (0.5μg/μl) were injected 3 times in left Tibialis Anterior (TA) of C57BL6/mdx mice: every 7 days for 21 days. The right TA was injected with vehicle (20 μL PBS1x) following the same timing described for EVs injection.

### RNA-sequencing

For RNA-sequencing sample preparation, MuSCs and FAPs were freshly isolated by FACS from 6 C57Bl6 mdx male mice 8-weeks old treated or not with TSA for 15 days. RNA was collected using Trizol reagent (#T9424, Sigma). About 100 ng/μL of total RNA was sent in duplicate to IGA (Istituto di Genomica Applicata, Udine) for RNA sequencing using Illumina TruSeq Stranded Total RNA kit Ribo-Zero GOLD on Illumina Hiseq2500 platform.

### Statistical Analysis

The number of independent experimental replications and precision measures are reported in the Figure legends (n, mean ± SEM).

Statistical analysis was performed using Prism 7.0 A software (Pad Software).

All data meet the assumptions of the tests (e.g., normal distribution). Unpaired, two-tailed Student’s *t* test were used to compare the means of two groups, while one-way analysis of variance (ANOVA) was used for comparison among the different groups while two-way analysis of variance was used for comparison among different groups to examines the influence of different independent variables on dependent variable. When ANOVA was significant, post hoc testing of differences between groups was performed using Tukey’s honestly significant difference (HSD).

A *P* < 0.05 was considered statistically significant.

No statistical method was used to predetermine sample size for animal studies. The animal experiments were not randomized. The investigators were not blinded to allocation during experiments and outcome assessment. No exclusion criteria were applied to exclude samples or animals from analysis.

## Data and Software availability

The cells positive for the stainings described in the text were quantified using ImageJ software (https://imagej.nih.gov/ij/download.html). The cross-sectional area (CSA) was also calculated using the ImageJ software and Macro seg 5 modif.ijm specific plugin. Fibrotic areas were measured from sections evaluating image analysis algorithms for colour deconvolution. ImageJ was used for image processing, the original image was segmented with three clusters and the plugin assumes images generated by color subtraction (white representing background, blue collagen, and magenta non collagen regions).

FACS profile analysis of MuSCs and FAPs were performed using Flowjo software (https://www.flowjo.com).

The RNA sequencing analysis was performed mapping more than 20 millions of reads for each sample to the Mus Musculus GRCm38.78 genome using TopHat 2.0.9. Read count was performed with HTSeq-0.6.1p1. Mapped reads were analysed with R-studio using DESeq2 to obtain normalized RPKM, P-Value, P-adjusted and log2fold changes values. Genes were considered differentially expressed if the 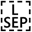 P-adjusted value was <0.1.

Additional raw data that support the findings of this study are available from the corresponding authors upon reasonable request. RNAseq data are available through SRA accession code SRP143532. miRNA pathway analysis was performed using miRPath based on predicted miRNA targets provided by the DIANA-microT-CDS algorithm and/or experimentally validated miRNA interactions derived from DIANA-TarBase v6.0 (http://www.microrna.gr/miRPathv2).

### Prediction of miRNA-mRNA interactions and network construction

Data from RNAseq (padj<0.1) in MuSCs (treated or not with TSA) and Microarray analysis in FAPs (treated or not with TSA) derived EVs, were uploaded in IPA (https://www.qiagenbioinformatics.com/products/ingenuity-pathway-analysis/). With the IPA tool “miR Target Filter” the miRNA upregulated in FAPs EVs were intersected with transcripts down-regulated in MuSCs after TSA treatment. Only interactions experimentally observed and highly predicted were selected for downstream analysis. *Notch* pathway was extrapolated from IPA database using the tool “grow”, while its modulation was predicted by the Molecular Activity Predictor (MAP) tool. The cartoon was realized with IPA path-designer.

## miR Oligo Table

|  |  |  |
| --- | --- | --- |
| MS00033740 | Qiagen | Hs_RNU6-2_1 |
| MS00001372 | Qiagen | Mm_miR-29a_1 |
| MS00033761 | Qiagen | Mm_miR-494_2 |
| MS00011704 | Qiagen | Mm_miR-30a_1 |
| MS00002387 | Qiagen | Mm_miR-449_1 |
| MS00001869 | Qiagen | Mm_miR206_1 |
| YCO0073796 | Qiagen | Mm_miR-145a-5p |

## Acknowledgments

We thank Dr Roberts and all members of Puri lab for reading the manuscript and for their insightful inputs. We thank them for critical discussion and daily support, we are grateful for precious comments during the experiments and manuscript preparation. We thank Dr. Mauro Manno - Institute of Biophysics (IBF), National Research Council (CNR) of Italy, Palermo, Italy for DLS assay.

We thank Giovanna Borsellino and Luca Battistini (Fondazione Santa Lucia, Roma, Italy) for isolation of MuSCs and FAPs by cell sorter.

This work has been supported by the following **funding**: Association Francaise contre les Myopathies, Italian Ministry of Health (GR-2016-02362451) to V:S:, MDA and EPIGEN F7 to P.L.P.DPP_Netherland post-Doc fellowship to V.S.; DPP_Ita PhD fellowship to L.T., AFM post-Doc fellowship to S.C.

## Authors Contribution

V.S. and M.S. performed most of the experimental work and contributed to experimental design, data analysis and discussions. In particular, M.S. performed the *in vivo* experiments, stainings on human and animal muscles sections. L.T. performed muscle fibre isolation and culture with FAP-derived EVs. M.Sc. performed SEM analysis. V.B. and A.B. contributed experimentally to EVs characterization. M.D.B. performed FACS sorting experiments and D.A. contributed to CytoFlex analysis of intramuscular injected EVs. S.C and L.T. performed bioinformatic data analysis. A.D.A. and E.B. provided human biopsies of Control and DMD patients. S.Ca and P.B were the sponsor of the clinical trial. M.S., S.C., V.S. and P.L.P conceived the experiments, supervised the study and interpreted the data. P.L.P. wrote the manuscript, and all authors discussed the results and commented on the manuscript.

## Data and materials availability

All data needed to evaluate the conclusions in the paper are present in the paper and/or the Supplementary Materials. Additional data related to this paper may be requested from the authors.

## Supplementary Materials

**Fig. S1:**
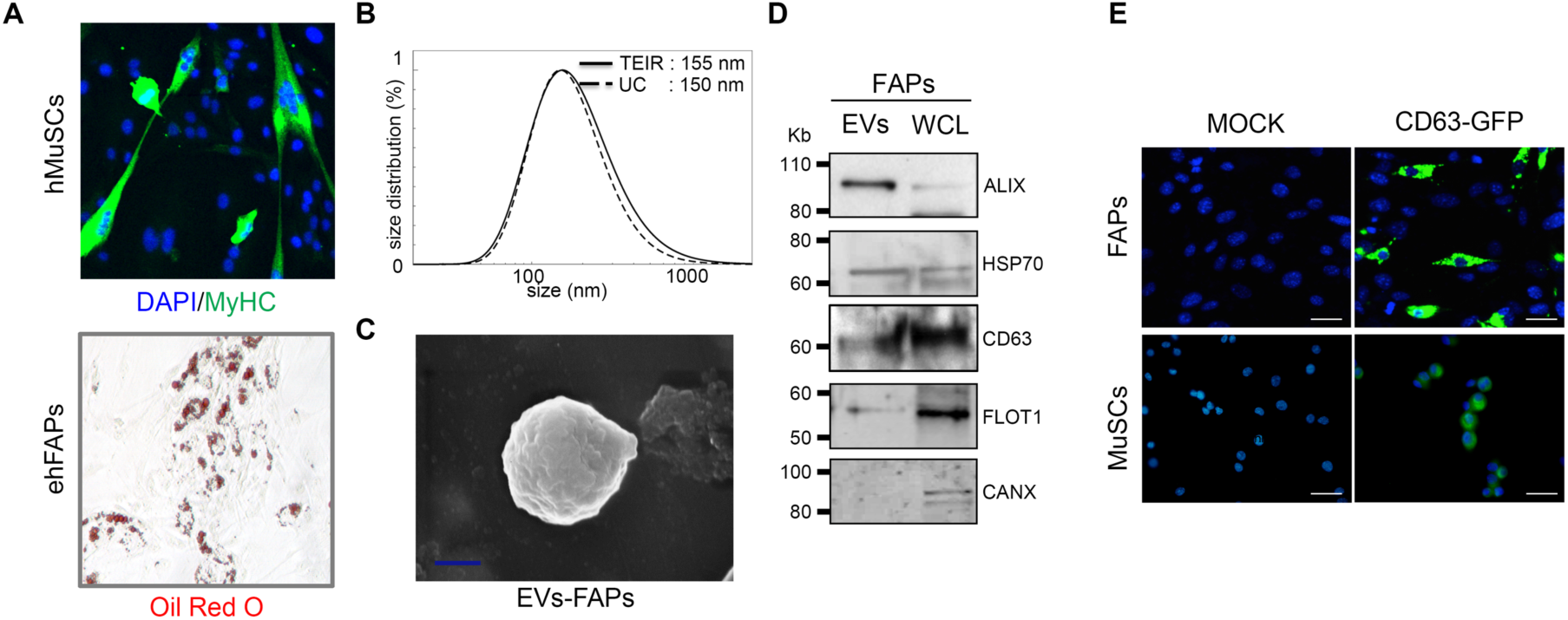
FAPs release Extracellular vesicles with physical features of exosomes that are transferred to MuSCs. **(A)** Representative images showing the differentiation of hMuSCs (MyHC staining-green – upper panel) and ehFAPs (Oil Red O staining-red – lower panel) isolated from biopsies of DMD patients before treatment with Giv. Dose and timing of Giv treatment as described in Bettica et al., 2016 (23). **(B)** Dynamic light scattering (DLS) analysis of FAPs-derived extracellular vesicles (EVs) isolated by total extracellular vesicles isolation reagent (TEIR) or ultracentrifugation (UC). **(C)** Scanned electron microscopy of FAPs-EV purified using TEIR. **(D)** Western blot analysis for ALIX, HSP70, CD63, Flotilin1 (FLOT1) and Calnexin (CANX) antibodies in FAPs-EVs and in the cell lysates (WCL). **(E)** Representative images of CD63-GFP detection (green) in MuSCs and FAPs co-cultured in transwell. Scale bar = 25 μm. Nuclei were counterstained with DAPI (blue).

**Fig. S2:**
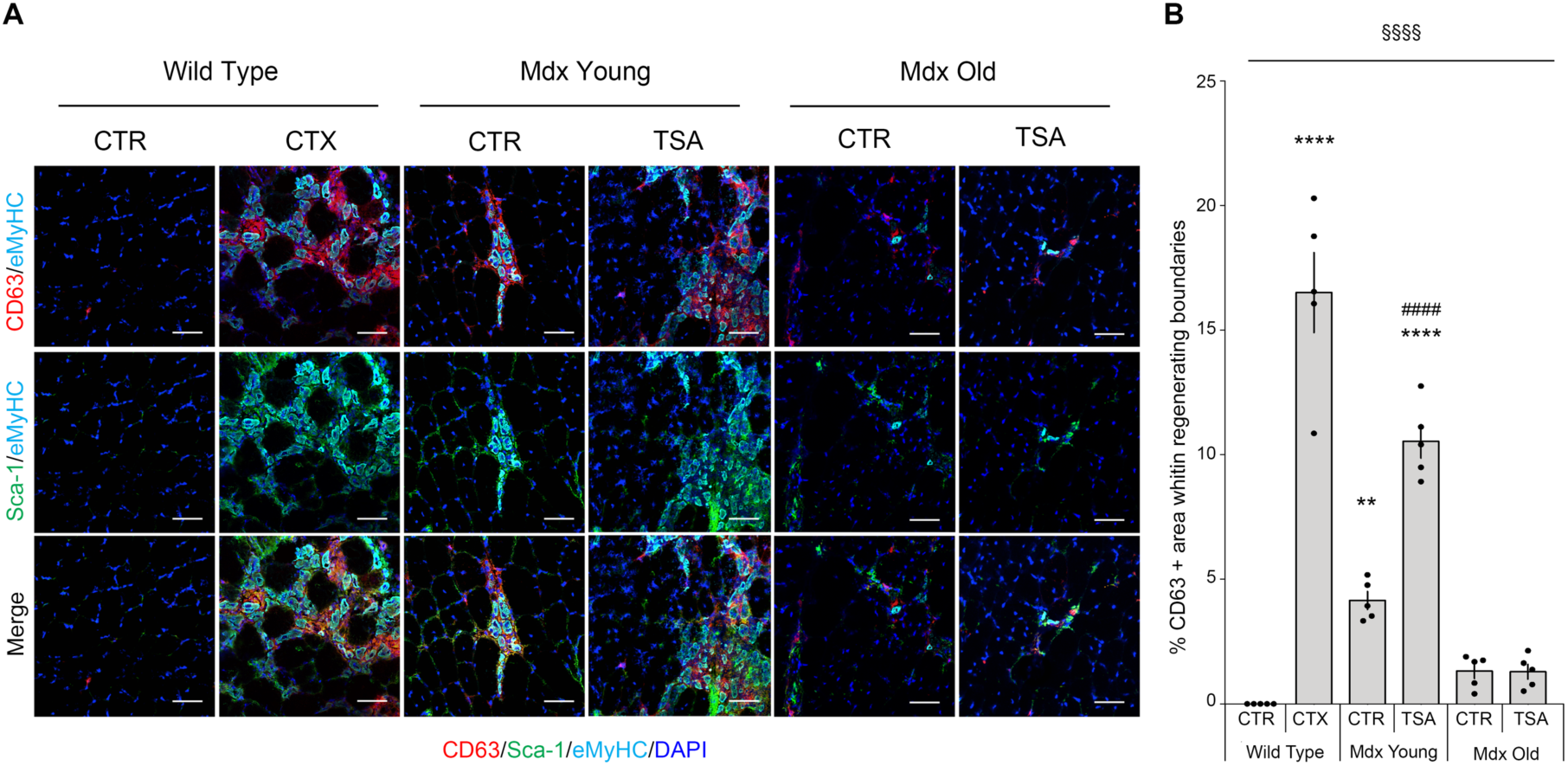
Increased number of EV associates with muscle regeneration. **(A)** Representative images of immunofluorescence for CD63 (red), Sca-1 (green) and eMyHC (cyan) in transversal sections of tibialis anterior muscles from: wild type mice unperturbed (control –CTR) or cardiotoxin injured (CTX); mdx young or old mice treated either with vehicle (control –CTR) or TSA (TSA, 0.6 mg/kg/die for 15 days by i.p injection). N=5. Scale bar = 50 μm. **(B)** Graph relative to the quantification of CD63 positive signal within the regenerative boundaries in the condition described in A. Star (*) indicates Tukey analysis compared to Wild Type CTR mice, **p < 0,01; ****p < 0,0001. Hash (#) means significance by Tukey analysis compared to Mdx Young CTR, #### p < 0,0001. § indicates Anova analysis, §§§§ p < 0,0001. Nuclei were counterstained with DAPI (blue). All data correspond to the average ± SEM.

**Fig. S3:**
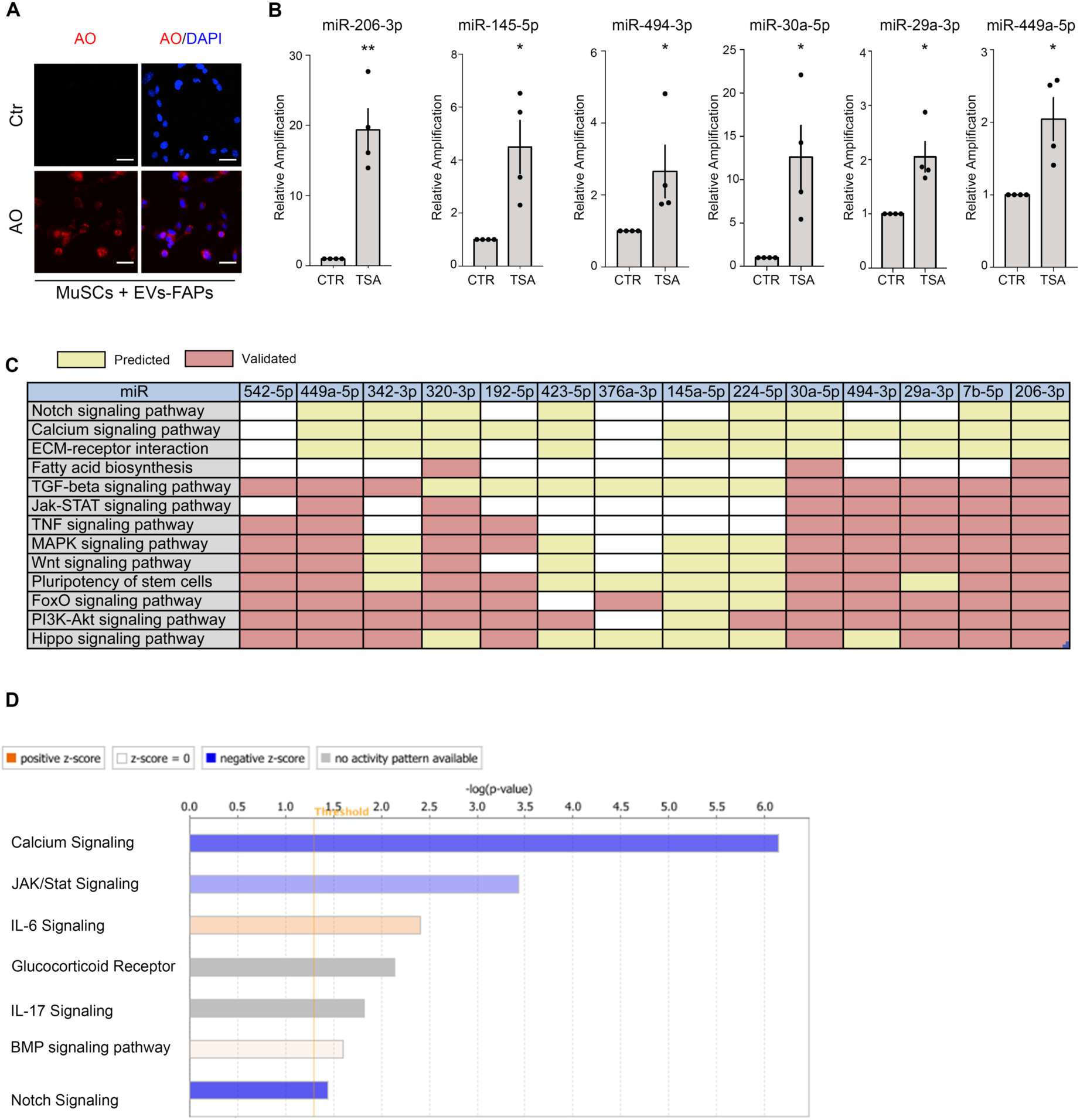
HDACi up-regulate a subset of miRs in FAP-derived EVs. **(A)** MuSCs uptake of RNA from FAPs-derived extracellular vesicles (EVs-FAPs). EVs harvested from FAPs donor cells were stained with Acridine Orange fluorescent dye (AO-red signal) or not (Ctr) and incubated with MuSCs. Nuclei were counterstained with DAPI (blue). Scale bar =50 μm. **(B)** Graphs showing the relative expression of microRNAs resulted up-regulated in EVs from FAPs by TSA treatment. Star (*) indicates statistical analysis by t-test relative to EVs CTR. *p <0.05, **p<0,01. All data correspond to the average ± SEM (n=4). **(C)** Table representing the Pathway Analysis of TSA up-regulated microRNAs induced in FAPs-derived EVs based on their predicted (shown in yellow) or already validated (in red) targets. **(D)** Ingenuity Pathway Analysis of canonical pathways found significantly modulated (activated (red) or suppressed (blue)) in MuSCs TSA compared with MuSCs CTR. The height of the bars reflect the −log(p value) for each pathway. Only pathways with −log(p value) > 1.3 are represented and considered statistically significant.

**Fig. S4:**
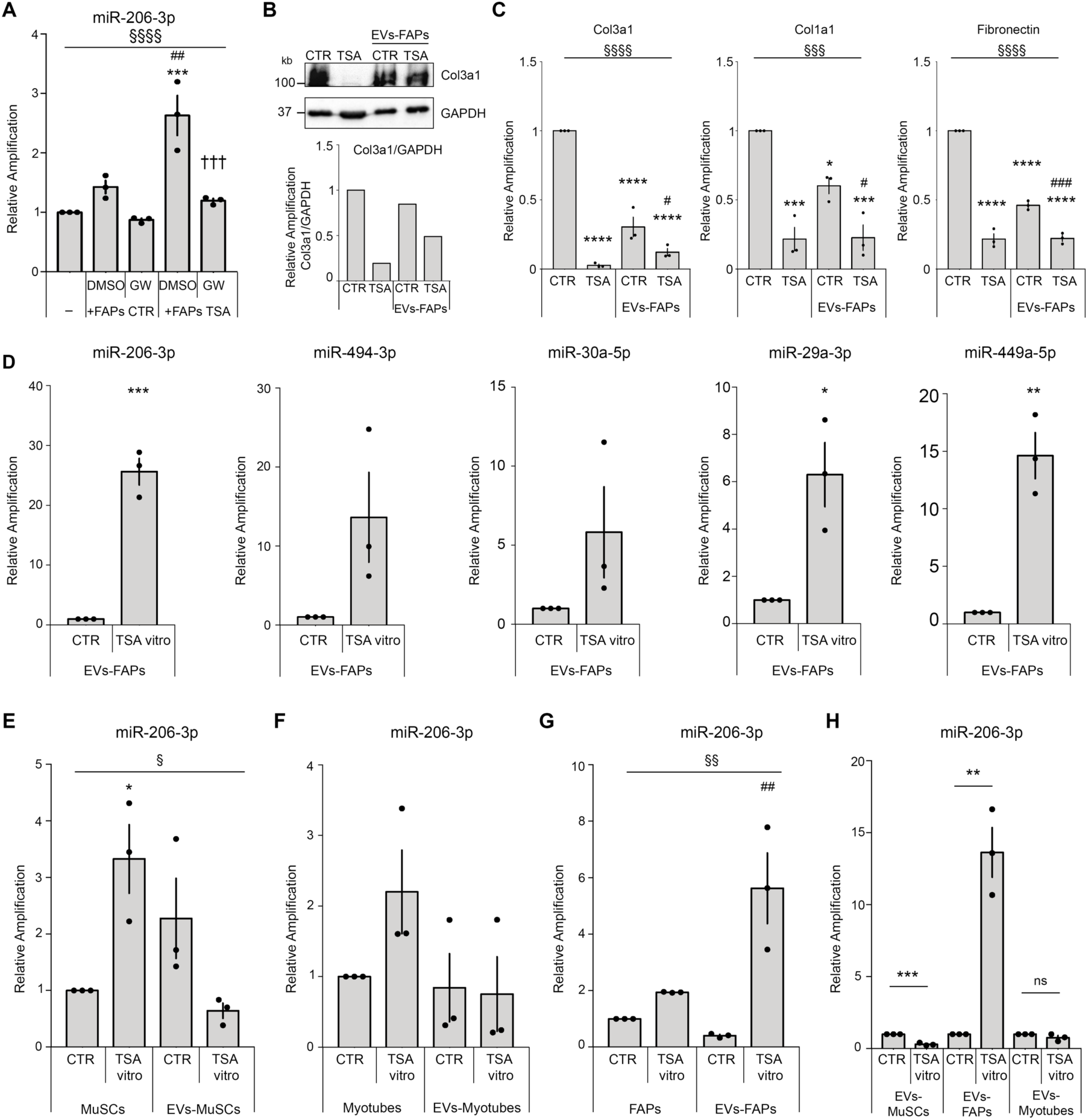
Direct and specific induction of miRs in EVs of FAPs by HDACi. **(A)** Graph showing the relative expression of miR-206-3p MuSCs cultured alone (-) or in co-colture with FAPs treated ex vivo with DMSO or with GW4869 (GW, 10 uM added to FAPs 30 minutes before the co-culture with MuSCs) isolated from mdx vehicle (+FAPs CTR) or TSA (+FAPs TSA) treated animals (n=3). Star (*) indicates statistical analysis by Tukey test relative to MuSCs alone (-); ***p<0.001. Hash (#) indicates statistical analysis by Tukey test relative to MuSCs in co-culture with FAPs CTR treated with DMSO; ^##^p<0.01. Cross (†) means Tukey analysis compared to MuSCs in co-culture with FAPs TSA treated with DMSO, ^†††^p<0,001. **(B)** Upper panel: Western Blot with Col3a1 and GAPDH antibodies using whole muscle extracts from mdx mice treated with vehicle (CTR) and TSA (TSA) or using lysates from FAP-derived EVs isolated from 1.5 old mdx mice treated with vehicle (EVs FAPs CTR) or TSA (EVs FAPs TSA). Lower panel: graph showing the quantification for the Western Blot in the upper panel, with Col3a1 relative to GAPDH. **(C)** Graphs showing the relative expression of Col3a1, Col1a1 and Fibronectin mRNA for the samples described in B (n=3). Star (*) indicates statistical analysis by Tukey test relative to CTR, ***p<0.001, ****p<0.0001. Hash (#) indicates statistical analysis by Tukey test relative to EVs FAPs CTR, ^#^p < 0.05. § represents statistical analysis by Anova test. §§§p < 0,001, §§§§p < 0,0001. **(D)** Graphs showing the relative expression of microRNAs resulted up-regulated in EVs isolated from FAPs after in vitro treatment with TSA (n=3). Star (*) means significance compared to CTR, *p<0.05; **p<0.01; ***p<0,001. **(E)** Graph showing miR206-3p relative expression into MuSCs treated in vitro with vehicle (CTR) or (TSA) and EVs from MuSCs (EVs-MuSCs) treated *in vitro* with vehicle (CTR) or TSA (TSA) (n=3). Star (*) indicates statistical analysis by Tukey test relative to MuSCs untreated (CTR), *p< 0.05. § indicates Anova analysis. §p<0,05. **(F)** Graph showing miR206-3p relative expression into myotubes (Myotubes) and myotubes derived EVs (EVs-Myotubes) treated *in vitro* with TSA (TSA vitro) or vehicle (CTR); (n=3). **(G)** Graph showing miR206-3p relative expression into FAPs (FAPs) and FAPs derived EVs (EVs-FAPs) treated *in vitro* with TSA (TSA vitro) or vehicle (CTR); (n=3). Hash (#) means significance compared to EVs-FAPs CTR, ##p<0,01. § indicates Anova analysis. §§p<0,01. **(H)** Graph showing miR206-3p relative expression into EVs derived from MuSCs, FAPs and Myotubes treated *in vitro* with TSA (TSA vitro) or vehicle (CTR). Star (*) indicates statistical analysis by t-test relative to the relative untreated (CTR) cell population, **p< 0.01, ***p<0.001, ns=not significant. TSA was administrated 0.6 mg/kg/day for 15 days. TSA in vitro was added to cells 50nM for 12hrs. All data correspond to the average ± SEM.

**Fig. S5:**
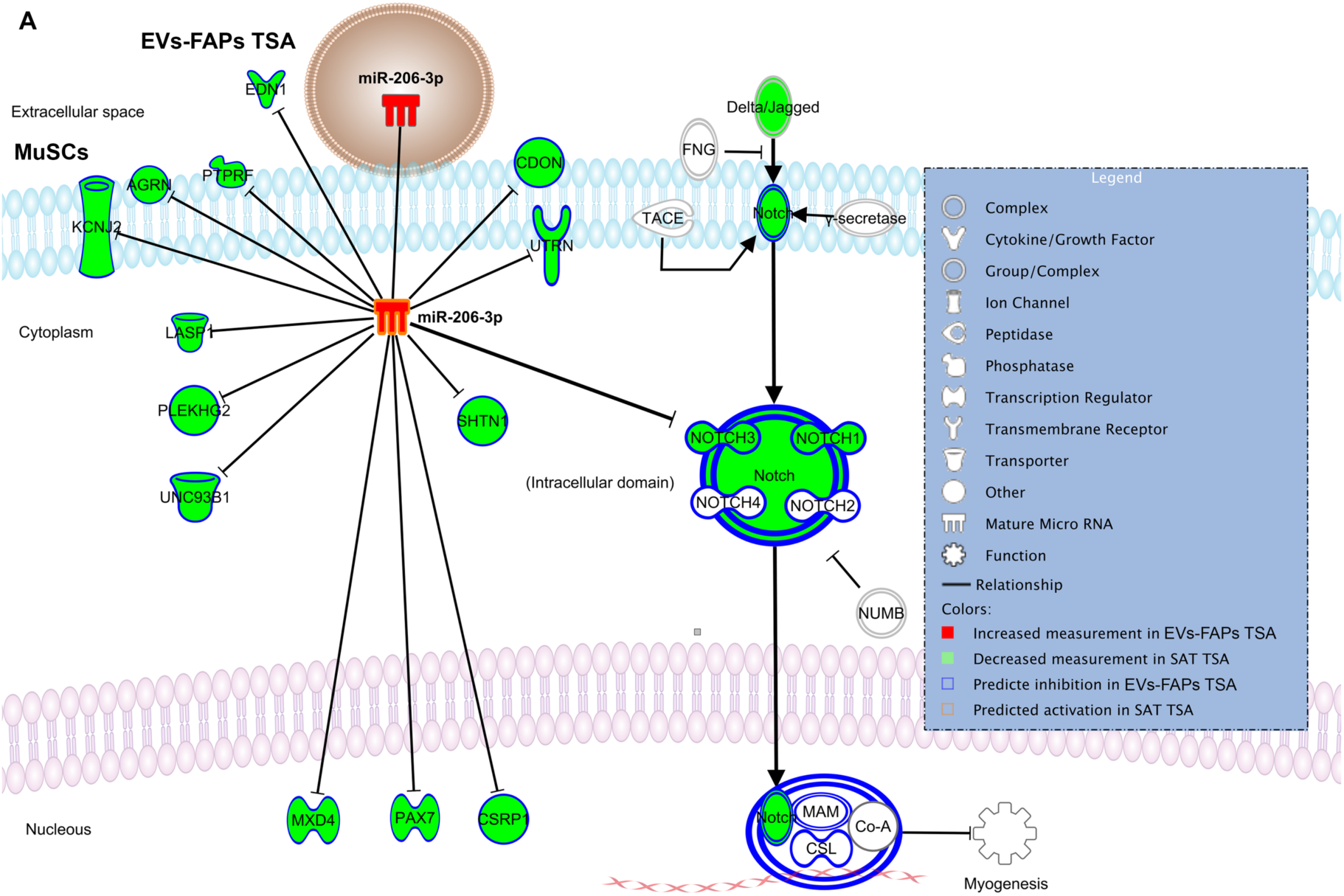
miR206 regulation of Notch3. Downstream analysis of putative miR206 targets annotated or predicted with high confidence. Ingenuity pathway analysis (IPA), including Notch3

## References

1. A. L. Moyer, K. R. Wagner, Regeneration versus fibrosis in skeletal muscle. Curr. Opin. Rheumatol. 23, 568–573 (2011).

2. J. G. Tidball, Mechanisms of muscle injury, repair, and regeneration. Compr. Physiol. 1, 2029–2062 (2011).

3. J. Farup, L. Madaro, P. L. Puri, U. R. Mikkelsen, Interactions between muscle stem cells, mesenchymal-derived cells and immune cells in muscle homeostasis, regeneration and disease. Cell Death Dis. 6, e1830 (2015).

4. A. Mauro, SATELLITE CELL OF SKELETAL MUSCLE FIBERS. J. Biophys. Biochem. Cytol. 9, 493–495 (1961).

5. C. F. Bentzinger, Y. X. Wang, N. A. Dumont, M. A. Rudnicki, Cellular dynamics in the muscle satellite cell niche. EMBO Rep. 14, 1062–1072 (2013).

6. N. A. Dumont, Y. X. Wang, M. A. Rudnicki, Intrinsic and extrinsic mechanisms regulating satellite cell function. Dev. Camb. Engl. 142, 1572–1581 (2015).

7. J. G. Tidball, K. Dorshkind, M. Wehling-Henricks, Shared signaling systems in myeloid cell-mediated muscle regeneration. Dev. Camb. Engl. 141, 1184–1196 (2014).

8. Y. Kharraz, J. Guerra, C. J. Mann, A. L. Serrano, P. Muñoz-Cánoves, Macrophage plasticity and the role of inflammation in skeletal muscle repair. Mediators Inflamm. 2013, 491497 (2013).

9. A. W. B. Joe, L. Yi, A. Natarajan, F. Le Grand, L. So, J. Wang, M. A. Rudnicki, F. M. V. Rossi, Muscle injury activates resident fibro/adipogenic progenitors that facilitate myogenesis. Nat. Cell Biol. 12, 153–163 (2010).

10. A. Uezumi, S. Fukada, N. Yamamoto, S. Takeda, K. Tsuchida, Mesenchymal progenitors distinct from satellite cells contribute to ectopic fat cell formation in skeletal muscle. Nat. Cell Biol. 12, 143–152 (2010).

11. B. Malecova, S. Gatto, U. Etxaniz, M. Passafaro, A. Cortez, C. Nicoletti, L. Giordani, A. Torcinaro, M. De Bardi, S. Bicciato, F. De Santa, L. Madaro, P. L. Puri, Dynamics of cellular states of fibro-adipogenic progenitors during myogenesis and muscular dystrophy. Nat. Commun. 9, 3670 (2018).

12. H. M. Blau, B. D. Cosgrove, A. T. V. Ho, The central role of muscle stem cells in regenerative failure with aging. Nat. Med. 21, 854–862 (2015).

13. A. Uezumi, T. Ito, D. Morikawa, N. Shimizu, T. Yoneda, M. Segawa, M. Yamaguchi, R. Ogawa, M. M. Matev, Y. Miyagoe-Suzuki, S. Takeda, K. Tsujikawa, K. Tsuchida, H. Yamamoto, S. Fukada, Fibrosis and adipogenesis originate from a common mesenchymal progenitor in skeletal muscle. J. Cell Sci. 124, 3654–3664 (2011).

14. D. R. Lemos, F. Babaeijandaghi, M. Low, C.-K. Chang, S. T. Lee, D. Fiore, R.-H. Zhang, A. Natarajan, S. A. Nedospasov, F. M. V. Rossi, Nilotinib reduces muscle fibrosis in chronic muscle injury by promoting TNF-mediated apoptosis of fibro/adipogenic progenitors. Nat. Med. 21, 786–794 (2015).

15. J. E. Heredia, L. Mukundan, F. M. Chen, A. A. Mueller, R. C. Deo, R. M. Locksley, T. A. Rando, A. Chawla, Type 2 innate signals stimulate fibro/adipogenic progenitors to facilitate muscle regeneration. Cell. 153, 376–388 (2013).

16. G. C. Minetti, C. Colussi, R. Adami, C. Serra, C. Mozzetta, V. Parente, S. Fortuni, S. Straino, M. Sampaolesi, M. Di Padova, B. Illi, P. Gallinari, C. Steinkühler, M. C. Capogrossi, V. Sartorelli, R. Bottinelli, C. Gaetano, P. L. Puri, Functional and morphological recovery of dystrophic muscles in mice treated with deacetylase inhibitors. Nat. Med. 12, 1147–1150 (2006).

17. S. Consalvi, V. Saccone, L. Giordani, G. Minetti, C. Mozzetta, P. L. Puri, Histone deacetylase inhibitors in the treatment of muscular dystrophies: epigenetic drugs for genetic diseases. Mol. Med. Camb. Mass. 17, 457–465 (2011).

18. C. Mozzetta, S. Consalvi, V. Saccone, M. Tierney, A. Diamantini, K. J. Mitchell, G. Marazzi, G. Borsellino, L. Battistini, D. Sassoon, A. Sacco, P. L. Puri, Fibroadipogenic progenitors mediate the ability of HDAC inhibitors to promote regeneration in dystrophic muscles of young, but not old Mdx mice. EMBO Mol. Med. 5, 626–639 (2013).

19. S. Consalvi, C. Mozzetta, P. Bettica, M. Germani, F. Fiorentini, F. Del Bene, M. Rocchetti, F. Leoni, V. Monzani, P. Mascagni, P. L. Puri, V. Saccone, Preclinical studies in the mdx mouse model of duchenne muscular dystrophy with the histone deacetylase inhibitor givinostat. Mol. Med. Camb. Mass. 19, 79–87 (2013).

20. V. Saccone, S. Consalvi, L. Giordani, C. Mozzetta, I. Barozzi, M. Sandoná, T. Ryan, A. Rojas-Muñoz, L. Madaro, P. Fasanaro, G. Borsellino, M. De Bardi, G. Frigè, A. Termanini, X. Sun, J. Rossant, B. G. Bruneau, M. Mercola, S. Minucci, P. L. Puri, HDAC-regulated myomiRs control BAF60 variant exchange and direct the functional phenotype of fibro-adipogenic progenitors in dystrophic muscles. Genes Dev. 28, 841–857 (2014).

21. Consalvi, Sandona, M, Saccone V, Epigenetic Reprogramming of Muscle Progenitors: Inspiration for Clinical Therapies. doi: 10.1155/2016/6093601 2016, 1–11 (2016).

22. M. Sandoná, S. Consalvi, L. Tucciarone, P. L. Puri, V. Saccone, HDAC inhibitors for muscular dystrophies: progress and prospects. Expert Opin. Orphan Drugs. 4, 125–127 (2016).

23. P. Bettica, S. Petrini, V. D’Oria, A. D’Amico, M. Catteruccia, M. Pane, S. Sivo, F. Magri, S. Brajkovic, S. Messina, G. L. Vita, B. Gatti, M. Moggio, P. L. Puri, M. Rocchetti, G. De Nicolao, G. Vita, G. P. Comi, E. Bertini, E. Mercuri, Histological effects of givinostat in boys with Duchenne muscular dystrophy. Neuromuscul. Disord. NMD. 26, 643–649 (2016).

24. S. Consalvi, V. Saccone, C. Mozzetta, Histone deacetylase inhibitors: a potential epigenetic treatment for Duchenne muscular dystrophy. Epigenomics. 6, 547–560 (2014).

25. K. Trajkovic, C. Hsu, S. Chiantia, L. Rajendran, D. Wenzel, F. Wieland, P. Schwille, B. Brügger, M. Simons, Ceramide triggers budding of exosome vesicles into multivesicular endosomes. Science. 319, 1244–1247 (2008).

26. C. Théry, S. Amigorena, G. Raposo, A. Clayton, Isolation and Characterization of Exosomes from Cell Culture Supernatants and Biological Fluids. Curr. Protoc. Cell Biol., 30(**1**), 3.22.1–3.22.29 (2006).

27. J. Van Deun, P. Mestdagh, R. Sormunen, V. Cocquyt, K. Vermaelen, J. Vandesompele, M. Bracke, O. De Wever, A. Hendrix, The impact of disparate isolation methods for extracellular vesicles on downstream RNA profiling. J. Extracell. Vesicles. 3 (2014), doi:10.3402/jev.v3.24858.

28. G. Raposo, W. Stoorvogel, Extracellular vesicles: Exosomes, microvesicles, and friends. J Cell Biol. 200, 373–383 (2013).

29. V. Sokolova, A.-K. Ludwig, S. Hornung, O. Rotan, P. A. Horn, M. Epple, B. Giebel, Characterisation of exosomes derived from human cells by nanoparticle tracking analysis and scanning electron microscopy. Colloids Surf. B Biointerfaces. 87, 146–150 (2011).

30. EV-TRACK Consortium, J. Van Deun, P. Mestdagh, P. Agostinis, Ö. Akay, S. Anand, J. Anckaert, Z. A. Martinez, T. Baetens, E. Beghein, L. Bertier, G. Berx, J. Boere, S. Boukouris, M. Bremer, D. Buschmann, J. B. Byrd, C. Casert, L. Cheng, A. Cmoch, D. Daveloose, E. De Smedt, S. Demirsoy, V. Depoorter, B. Dhondt, T. A. P. Driedonks, A. Dudek, A. Elsharawy, I. Floris, A. D. Foers, K. Gärtner, A. D. Garg, E. Geeurickx, J. Gettemans, F. Ghazavi, B. Giebel, T. G. Kormelink, G. Hancock, H. Helsmoortel, A. F. Hill, V. Hyenne, H. Kalra, D. Kim, J. Kowal, S. Kraemer, P. Leidinger, C. Leonelli, Y. Liang, L. Lippens, S. Liu, A. Lo Cicero, S. Martin, S. Mathivanan, P. Mathiyalagan, T. Matusek, G. Milani, M. Monguió-Tortajada, L. M. Mus, D. C. Muth, A. Németh, E. N. M. Nolte-’t Hoen, L. O’Driscoll, R. Palmulli, M. W. Pfaffl, B. Primdal-Bengtson, E. Romano, Q. Rousseau, S. Sahoo, N. Sampaio, M. Samuel, B. Scicluna, B. Soen, A. Steels, J. V. Swinnen, M. Takatalo, S. Thaminy, C. Théry, J. Tulkens, I. Van Audenhove, S. van der Grein, A. Van Goethem, M. J. van Herwijnen, G. Van Niel, N. Van Roy, A. R. Van Vliet, N. Vandamme, S. Vanhauwaert, G. Vergauwen, F. Verweij, A. Wallaert, M. Wauben, K. W. Witwer, M. I. Zonneveld, O. De Wever, J. Vandesompele, A. Hendrix, EV-TRACK: transparent reporting and centralizing knowledge in extracellular vesicle research. Nat. Methods. 14, 228–232 (2017).

31. H. Valadi, K. Ekström, A. Bossios, M. Sjöstrand, J. J. Lee, J. O. Lötvall, Exosome-mediated transfer of mRNAs and microRNAs is a novel mechanism of genetic exchange between cells. Nat. Cell Biol. 9, 654–659 (2007).

32. F. Collino, M. C. Deregibus, S. Bruno, L. Sterpone, G. Aghemo, L. Viltono, C. Tetta, G. Camussi, Microvesicles derived from adult human bone marrow and tissue specific mesenchymal stem cells shuttle selected pattern of miRNAs. PloS One. 5, e11803 (2010).

33. A. Montecalvo, A. T. Larregina, W. J. Shufesky, D. B. Stolz, M. L. G. Sullivan, J. M. Karlsson, C. J. Baty, G. A. Gibson, G. Erdos, Z. Wang, J. Milosevic, O. A. Tkacheva, S. J. Divito, R. Jordan, J. Lyons-Weiler, S. C. Watkins, A. E. Morelli, Mechanism of transfer of functional microRNAs between mouse dendritic cells via exosomes. Blood. 119, 756–766 (2012).

34. A. V. Vlassov, S. Magdaleno, R. Setterquist, R. Conrad, Exosomes: current knowledge of their composition, biological functions, and diagnostic and therapeutic potentials. Biochim. Biophys. Acta. 1820, 940–948 (2012).

35. Y. Nakamura, S. Miyaki, H. Ishitobi, S. Matsuyama, T. Nakasa, N. Kamei, T. Akimoto, Y. Higashi, M. Ochi, Mesenchymal-stem-cell-derived exosomes accelerate skeletal muscle regeneration. FEBS Lett. 589, 1257–1265 (2015).

36. C. S. Fry, T. J. Kirby, K. Kosmac, J. J. McCarthy, C. A. Peterson, Myogenic Progenitor Cells Control Extracellular Matrix Production by Fibroblasts during Skeletal Muscle Hypertrophy. Cell Stem Cell. 20, 56–69 (2017).

37. S. Greco, M. De Simone, C. Colussi, G. Zaccagnini, P. Fasanaro, M. Pescatori, R. Cardani, R. Perbellini, E. Isaia, P. Sale, G. Meola, M. C. Capogrossi, C. Gaetano, F. Martelli, Common micro-RNA signature in skeletal muscle damage and regeneration induced by Duchenne muscular dystrophy and acute ischemia. FASEB J. Off. Publ. Fed. Am. Soc. Exp. Biol. 23, 3335–3346 (2009).

38. X. H. Wang, MicroRNA in myogenesis and muscle atrophy. Curr. Opin. Clin. Nutr. Metab. Care. 16, 258–266 (2013).

39. Q. Zhao, Y. Kang, H.-Y. Wang, W.-J. Guan, X.-C. Li, L. Jiang, X.-H. He, Y.-B. Pu, J.-L. Han, Y.-H. Ma, Q.-J. Zhao, Expression profiling and functional characterization of miR-192 throughout sheep skeletal muscle development. Sci. Rep. 6, 30281 (2016).

40. M. G. Guess, K. K. B. Barthel, B. C. Harrison, L. A. Leinwand, miR-30 family microRNAs regulate myogenic differentiation and provide negative feedback on the microRNA pathway. PloS One. 10, e0118229 (2015).

41. H. Yamamoto, K. Morino, Y. Nishio, S. Ugi, T. Yoshizaki, A. Kashiwagi, H. Maegawa, MicroRNA-494 regulates mitochondrial biogenesis in skeletal muscle through mitochondrial transcription factor A and Forkhead box j3. Am. J. Physiol. Endocrinol. Metab. 303, E1419–1427 (2012).

42. L. Wang, L. Zhou, P. Jiang, L. Lu, X. Chen, H. Lan, D. C. Guttridge, H. Sun, H. Wang, Loss of miR-29 in myoblasts contributes to dystrophic muscle pathogenesis. Mol. Ther. J. Am. Soc. Gene Ther. 20, 1222–1233 (2012).

43. N. Liu, A. H. Williams, J. M. Maxeiner, S. Bezprozvannaya, J. M. Shelton, J. A. Richardson, R. Bassel-Duby, E. N. Olson, microRNA-206 promotes skeletal muscle regeneration and delays progression of Duchenne muscular dystrophy in mice. J. Clin. Invest. 122, 2054–2065 (2012).

44. G. Ma, Y. Wang, Y. Li, L. Cui, Y. Zhao, B. Zhao, K. Li, MiR-206, a Key Modulator of Skeletal Muscle Development and Disease. Int. J. Biol. Sci. 11, 345–352 (2015).

45. D. Cacchiarelli, J. Martone, E. Girardi, M. Cesana, T. Incitti, M. Morlando, C. Nicoletti, T. Santini, O. Sthandier, L. Barberi, A. Auricchio, A. Musarò, I. Bozzoni, MicroRNAs involved in molecular circuitries relevant for the Duchenne muscular dystrophy pathogenesis are controlled by the dystrophin/nNOS pathway. Cell Metab. 12, 341–351 (2010).

46. A. H. Williams, G. Valdez, V. Moresi, X. Qi, J. McAnally, J. L. Elliott, R. Bassel-Duby, J. R. Sanes, E. N. Olson, MicroRNA-206 delays ALS progression and promotes regeneration of neuromuscular synapses in mice. Science. 326, 1549–1554 (2009).

47. H. Aswad, A. Forterre, O. P. B. Wiklander, G. Vial, E. Danty-Berger, A. Jalabert, A. Lamazière, E. Meugnier, S. Pesenti, C. Ott, K. Chikh, S. El-Andaloussi, H. Vidal, E. Lefai, J. Rieusset, S. Rome, Exosomes participate in the alteration of muscle homeostasis during lipid-induced insulin resistance in mice. Diabetologia. 57, 2155–2164 (2014).

48. M. I. Rosenberg, S. A. Georges, A. Asawachaicharn, E. Analau, S. J. Tapscott, MyoD inhibits Fstl1 and Utrn expression by inducing transcription of miR-206. J. Cell Biol. 175, 77–85 (2006).

49. A. Amirouche, H. Tadesse, P. Miura, G. Bélanger, J. A. Lunde, J. Côté, B. J. Jasmin, Converging pathways involving microRNA-206 and the RNA-binding protein KSRP control post-transcriptionally utrophin A expression in skeletal muscle. Nucleic Acids Res. 42, 3982–3997 (2014).

50. J.-F. Chen, Y. Tao, J. Li, Z. Deng, Z. Yan, X. Xiao, D.-Z. Wang, microRNA-1 and microRNA-206 regulate skeletal muscle satellite cell proliferation and differentiation by repressing Pax7. J. Cell Biol. 190, 867–879 (2010).

51. J. Gagan, B. K. Dey, R. Layer, Z. Yan, A. Dutta, Notch3 and Mef2c proteins are mutually antagonistic via Mkp1 protein and miR-1/206 microRNAs in differentiating myoblasts. J. Biol. Chem. 287, 40360–40370 (2012).

52. T. Kitamoto, K. Hanaoka, Notch3 null mutation in mice causes muscle hyperplasia by repetitive muscle regeneration. Stem Cells Dayt. Ohio. 28, 2205–2216 (2010).

53. P. S. Zammit, J. P. Golding, Y. Nagata, V. Hudon, T. A. Partridge, J. R. Beauchamp, Muscle satellite cells adopt divergent fates. J. Cell Biol. 166, 347–357 (2004).

54. Y. Lee, S. El Andaloussi, M. J. A. Wood, Exosomes and microvesicles: extracellular vesicles for genetic information transfer and gene therapy. Hum. Mol. Genet. 21, R125–134 (2012).

55. N. Iraci, T. Leonardi, F. Gessler, B. Vega, S. Pluchino, Focus on Extracellular Vesicles: Physiological Role and Signalling Properties of Extracellular Membrane Vesicles. Int. J. Mol. Sci. 17, 171 (2016).

56. J. S. Choi, H. I. Yoon, K. S. Lee, Y. C. Choi, S. H. Yang, I.-S. Kim, Y. W. Cho, Exosomes from differentiating human skeletal muscle cells trigger myogenesis of stem cells and provide biochemical cues for skeletal muscle regeneration. J. Control. Release Off. J. Control. Release Soc. 222, 107–115 (2016).

57. N. A. Dumont, Y. X. Wang, J. von Maltzahn, A. Pasut, C. F. Bentzinger, C. E. Brun, M. A. Rudnicki, Dystrophin expression in muscle stem cells regulates their polarity and asymmetric division. Nat. Med. 21, 1455–1463 (2015).

58. N. C. Chang, F. P. Chevalier, M. A. Rudnicki, Satellite Cells in Muscular Dystrophy - Lost in Polarity. Trends Mol. Med. 22, 479–496 (2016).

59. C. Colussi, C. Mozzetta, A. Gurtner, B. Illi, J. Rosati, S. Straino, G. Ragone, M. Pescatori, G. Zaccagnini, A. Antonini, G. Minetti, F. Martelli, G. Piaggio, P. Gallinari, C. Steinkuhler, C. Steinkulher, E. Clementi, C. Dell’Aversana, L. Altucci, A. Mai, M. C. Capogrossi, P. L. Puri, C. Gaetano, HDAC2 blockade by nitric oxide and histone deacetylase inhibitors reveals a common target in Duchenne muscular dystrophy treatment. Proc. Natl. Acad. Sci. U. S. A. 105, 19183–19187 (2008).

60. L. Tucciarone, U. Etxaniz, M. Sandoná, S. Consalvi, P. L. Puri, V. Saccone, in Duchenne Muscular Dystrophy, C. Bernardini, Ed. (Springer New York, New York, NY, 2018; http://link.springer.com/10.1007/978-1-4939-7374-3_17), vol. 1687, pp. 231–256 (2018).

61. R. Noto, M. G. Santangelo, S. Ricagno, M. R. Mangione, M. Levantino, M. Pezzullo, V. Martorana, A. Cupane, M. Bolognesi, M. Manno, The tempered polymerization of human neuroserpin. PloS One. 7, e32444 (2012).

62. M. Logozzi, D. F. Angelini, E. Iessi, D. Mizzoni, R. Di Raimo, C. Federici, L. Lugini, G. Borsellino, A. Gentilucci, F. Pierella, V. Marzio, A. Sciarra, L. Battistini, S. Fais, Increased PSA expression on prostate cancer exosomes in in vitro condition and in cancer patients. Cancer Lett. 403, 318–329 (2017).

